# Profound immunomodulatory effects of ^225^Ac-NM600 drive enhanced anti-tumor response in prostate cancer

**DOI:** 10.1101/2022.09.26.509374

**Authors:** Carolina A. Ferreira, Hemanth K. Potluri, Christopher Massey, Joseph J. Grudzinski, Amanda Carston, Nathan Clemons, Anna Thickens, Zachary Rosenkrans, Cynthia Choi, Anatoly Pinchuk, Ohyun Kwon, Justin J. Jeffery, Bryan Bednarz, Zachary Morris, Jamey Weichert, Douglas G. McNeel, Reinier Hernandez

## Abstract

An immunosuppressive tumor microenvironment has hampered the efficacy of immunotherapy in prostate cancer. However, radiation-induced immunological effects can partly mediate anti-tumor effects by promoting a pro-inflammatory environment potentially responsive to immunotherapy. Herein, we examined the immunomodulatory properties of a radiopharmaceutical therapy (RPT) with NM600 radiolabeled with either a beta or alpha emitter in two prostate cancer models. ^225^Ac-NM600, but not ^177^Lu-NM600, promoted significant anti-tumor effects and improved overall survival. Immunomodulatory effects were dose, radionuclide, and tumor type-dependent. ^225^Ac-NM600 elicited an array of immunomodulatory effects such as increased CD8/Treg ratio, activation of effector and memory T cells, abrogation of infiltrating suppressor cells (e.g., Tregs and MDSCs), and increased levels of Th1 cytokine and pro-inflammatory chemokines. Importantly, we demonstrate the need to carefully characterize the immune responses elicited by RPT both pre-clinically and clinically to maximize tumor control and avoid potential counterproductive immunosuppressive effects.

**Teaser:** Targeted alpha therapy can create a pro-inflammatory tumor micro-environment that partly explains stronger anti-tumor responses in prostate cancer

## INTRODUCTION

Prostate cancer (PCa) still accounts for greater than 350,000 deaths worldwide per year. Due to its unique reliance on androgens for growth and progression, androgen deprivation constitutes the first line of therapy for metastatic prostate cancer. Unfortunately, despite castration levels of serum testosterone, most patients eventually develop metastatic castration-resistant prostate cancer (mCRPC), whose mortality rate exceeds 50% with a median life expectancy of fewer than three years (*1*). Bone metastasis occurs in 90% of men with mCRPC and is associated with a poor prognosis and diminished quality of life due to debilitating morbidities such as pain, bone fractures, and bone marrow failure (*2, 3*). Significant advances in the understanding of prostate cancer biology have led to the development and approval of several new agents in mCRPC (*1, 4*). However, over the last decade, these treatments have only contributed to a 5% decrease in mortality of mCRPC (*5*). These figures highlight the need for novel therapeutic combinations and innovative targeted therapies that can significantly improve survival over existing treatments.

For over half a century, external beam radiotherapy (EBRT) has been a pillar in the curative treatment of non-metastatic prostate cancer, with equivalent efficacy to radical prostatectomy (*6*). However, because treatments with large radiation fields can severely damage healthy tissue, EBRT is constrained in the number and size of tumor lesions it can treat in the latter stages of disease (*7*). Accordingly, in mCRPC, EBRT is only offered palliatively to alleviate acute pain due to bone metastases (*8*). Radiopharmaceutical therapy (RPT), however, involves systemic delivery of radioactive atoms to induce DNA damage to tumor cells selectively and can, therefore, reach multiple tumor sites at once while minimizing off-target tissue toxicity (*9*). Non-prostate-cancer-specific systemic radiation therapy approaches, such as Strontium-89 chloride (Metastron), Samarium-153 (Quadramet), and Radium-223 dichloride (Xofigo), are already clinically used to treat mCRPC that spreads to the bones (*10*). However, the FDA recently approved the first RPT agent (^177^Lu-PSMA-617) to treat mCRPC patients expressing the prostate-specific membrane antigen (PSMA) and that failed androgen receptor inhibition and taxane-based chemotherapy (*11*). ^177^Lu-PSMA-617 showed favorable dosimetry and encouraging response in terms of both biochemical — reduction of PSA levels by half — and radiographic responses. While this certainly represents a significant advancement in mCRPC management, ^177^Lu-PSMA-617 only extends survival a modest 4 months. In addition, around a third of patients never respond and toxicity is a concern (*12*). Development of new RPT agents as single agents and in combination with other synergistic therapies is warranted.

Recently explored strategies to improve RPT efficacy include using alpha particle emitting radionuclides (e.g., ^225^Ac, ^211^At, and ^212^Pb) instead of conventional beta-emitting isotopes. Alpha-particles, due to their significantly higher energies than beta particles (4-9 MeV versus 0.1 – 2.2 MeV) and shorter path length, have a high linear energy transfer (LET) with increased direct DNA double-strand breaks and enhanced cell kill, even in hypoxic conditions (*13–15*). Given the dosimetric advantages of alpha emitters compared to beta emitters, an increasing number of alpha-RPT agents have been tested in prostate cancer with encouraging results, including several PSMA ligands radiolabeled with ^225^Ac (*12*), ^212^Pb (*16*), and ^213^Bi (*17*), (*18*). However, few systematic, dosimetry-based studies have evaluated and compared the radiobiological effects of RPT with beta and alpha-emitters, and even fewer have been performed in the context of a fully functional immune system.

Our group has exploited the capacity of cancer cells to selectively sequester and retain metabolically resistant phospholipid (*19*) to develop a series of alkylphosphocholine (APC) chelate analogs targeting a broad spectrum of malignancies, including prostate cancer (*20*). NM600, an APC analog featuring a DOTA chelating moiety, shows favorable tumor targeting and pharmacokinetics (*21–23*) and can be labeled with various radiometals for imaging and RPT, including ^86^Y, ^90^Y, ^177^Lu, ^64^Cu, and ^225^Ac. Treatment with ^177^Lu-NM600 and ^90^Y-NM600 resulted in significant tumor growth inhibition and extended survival in multiple murine cancer models (*21–23*). We also demonstrated that these agents could elicit immunomodulatory effects on the tumor microenvironment (TME), enhancing immune susceptibility and anti-tumor response to immunotherapy. Notably, these effects manifested at relatively low absorbed radiation doses that did not elicit significant therapeutic responses. Interestingly, in a recent study (*20*), we demonstrated that ^90^Y-NM600 treatment led to the enrichment of immunosuppressive regulatory T cells (T_reg_) in the TME and poor efficacy in combination with anti-PD1 immune-check point inhibition in Tramp-C1 and MyC-CaP murine models of prostate cancer.

As the radiobiological and immunomodulatory effects of RPT are dose, fractionation, and radiation type dependent (*24, 25*), herein we set out to investigate and compare (1) the immunological effects on prostate tumor cells and associated TME and (2) anti-tumor efficacy of NM600 radiolabeled with the β-emitter ^177^Lu and the α-emitter ^225^Ac. We have implemented a theranostic approach employing SPECT/CT imaging to describe the distribution and estimate the dosimetry of ^177^Lu/^225^Ac-NM600. This enabled the systematic comparison between ^177^Lu-NM600 and ^225^Ac-NM600 on the magnitude and timing of the anti-tumor and immunomodulatory effects in two immunocompetent murine prostate cancer models. Our results point to significant advantages in terms of anti-tumor efficacy of ^225^Ac-NM600 over its β-emitting congeners, stemming from the marked ability of α-particles to stimulate pro-inflammatory cytokine expression and abrogate immunosuppressive cell lineages within the TME. Importantly, these results provide a strong rationale for using ^225^Ac-NM600 in conjunction with immunotherapies in advanced prostate cancer.

## MATERIALS AND METHODS

### Radiochemistry

The radiolabeling of 2-(trimethylammonio)ethyl(18-(4-(2-(4,7,10-tris(carboxymethyl)-1,4,7,10-tetraazacyclododecan1-yl)acetamido)phenyl)octadecyl) phosphate (NM600) with ^177^Lu or ^225^Ac proceeded by mixing the radiometal with NM600 (10-100μg per mCi) in 0.1M NaOAc buffer (pH 5.5) for 30 min at 90°C. The labeled compounds were purified using a reverse-phase Waters Oasis HLB Light (Milford, MA), eluted using 100% ethanol, dried with a stream of N_2_, and reconstituted in 0.9% NaCl with 0.4% v/v Tween 20. Yield and purity were determined with instant thin-layer chromatography using 50mM EDTA and silica-impregnated paper (Perkin Elmer). A cyclone phosphor image reader was used to analyze the chromatograms; the labeled compound remained at the spotting point while free radiometals moved with the solvent front.

### Animal Models

Tramp-C1 (ATCC, CRL-2730), a transgenic C57BL/6 prostate adenocarcinoma cell line, and MyC-CaP (ATCC, CRL-3255), a murine FVB epithelial-like prostate cancer cell line, were both cultured in Dulbecco’s Modified Eagles Medium (DMEM) supplemented with 4 mM L-glutamine, 4500 mg/L glucose, 1 mM sodium pyruvate, and 1500 mg/L sodium bicarbonate, cells were kept at 5% CO_2_ and 37°C.

All animal experiments were performed under an approved University of Wisconsin Institutional Animal Care and Use Committee (IACUC) protocol. 6-week-old male C57BL/6 mice and 6-week-old male FVB/NJ were ordered from Jax. After acclimation for one week, Tramp-C1 cells were inoculated subcutaneously (s.q.) into the right flank of C57BL/6 mice at 1 x 10^6^ cells/100uL of sterile 1x PBS mixed 1:1 with Matrigel. MyC-CaP cells were inoculated s.q. into FVB/NJ mice at 1 x 10^6^ cells/100uL. Tumors were allowed to grow to a size of ~200 mm^3^ prior to animal use in imaging or therapy studies. Tramp-C1 and MyC-CaP tumor volumes were assessed 3 times weekly via digital caliper measurement. Tumor volume was calculated using the formula for ellipsoid volume 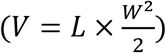 with L and W being the length and width of the tumor, respectively.

### SPECT/CT Imaging

Animals with subcutaneous MyC-Cap (FVB/NJ) or Tramp-C1 (C57BL/6) tumors were administered 3.7-7.4 MBq of ^177^Lu-NM600 in the lateral tail vein. Mice (n = 4) were placed prone into a MILabs U-SPECT^6^/CT^Uhr^ system (Houten, The Netherlands) under 2% isoflurane for longitudinal scans at 5, 24, 48, and 72 h post-injection (p.i.). CT scans were acquired for anatomical reference and attenuation correction and fused with the SPECT scan. Image reconstruction used a similarity-regulated ordered-subset expectation maximization (SROSEM) algorithm. Images were quantitatively analyzed by drawing volumes of interest (VOI) over the tumor and organs of interest to determine the percent injected activity (IA) per gram (%IA/g) for each tissue. *Ex vivo* biodistribution study was carried out after the last scan time point.

### *Ex Vivo* Biodistribution

Mice bearing either MyC-Cap (FVB/NJ) or Tramp-C1 (C57BL/6) subcutaneous were injected with 3.7 MBq of ^177^Lu-NM600, 3.7MBq of ^86^Y-NM600, or 7.4 KBq of ^225^Ac-NM600. Animals were euthanized via CO_2_ asphyxiation at 4, 24, 72, 120, and 192h (n = 3) post-injection, and organs of interest were collected for *ex vivo* biodistribution. Organs from animals that received ^225^Ac-NM600 were allowed to decay in storage overnight to reach secular equilibrium with ^213^Bi, which could be quantified. Organs were wet weighted, analyzed with a Perkin Elmer Wizard2 Gamma Counter (Westham, MA), and decay corrected to calculate the %IA/g for each tissue.

### Dosimetry Estimations

To estimate dosimetry of ^225^Ac-NM600, *ex vivo* biodistribution results were used. Allometric scaling was first performed to estimate the organ mass with respect to body mass for each mouse model, assuming 30 g for MyC-CaP and 25 g for TRAMP. Then, the residence time in each organ, 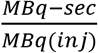, was determined using trapezoidal integration, assuming only physical decay after the last time point. Lastly, the residence time for each organ was multiplied by the dose factor, 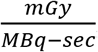, for each organ which was assumed to be a sphere of equal mass that only receives self-dose. The total dose for each organ was computed by summing all contributions from the complete decay chain of ^225^Ac, considering negligible redistribution of daughter isotopes. No correction for relative biological effectiveness (RBE = 1) was implemented on the dosimetry estimates.

SPECT/CT-based ^177^Lu-NM600 dosimetry was estimated according to a previously described method (*22, 26*). Total injected activity within each organ was calculated from extrapolation of %IA/g at a given time point. The standard mouse model was used to convert organ-specific cumulative activity into absorbed dose per injected activity (Gy/MBq). Dose contributions from surrounding organs were also included in calculations

### Toxicity Evaluations

To evaluate ^177^Lu-NM600 and ^225^Ac-NM600 toxicity, comprehensive metabolic panel (CMP), complete blood count (CBC), and histology studies were performed in control and therapy mice. Groups of MyC-Cap bearing (FVB/NJ) and Tramp-C1 bearing (C57BL/6) (n = 9) were administered 5.55 (low dose) or 18.5 MBq (high dose) of ^177^Lu-NM600, 7.4 (low dose) or 18.5 KBq (high dose) of ^225^Ac-NM600, or vehicle IV. Three mice were culled from each cohort on days 7, 14, and 28 p.i. and 500 μL of blood was collected via axillary bleed. CBC analysis was completed using whole blood and an Abaxis VetScan HM5 hematology analyzer (Union City, CA). The remaining blood was centrifuged at 4000 rpm for 10 minutes to separate the serum from red blood cells; the serum was then run on an Abaxis VetScan VS2 analyzer (Union City, CA). Toxicity tests were performed in control mice for baseline comparison. Organs with marked radiosensitivity or showing elevated ^177^Lu/^225^Ac-NM600 uptake, such as the liver, spleen, kidneys, femur, and small intestine, were collected and fixed for H&E staining.

### Therapeutic Studies

Mice with either MyC-Cap (FVB/NJ) or Tramp-C1 (C57BL/6) subcutaneous tumors were treated when tumors reached an approximate volume of 150 mm^3^. Mice were randomized into groups of n = 10 and injected with 5.55 (low dose) or 18.5 MBq (high dose) of ^177^Lu-NM600, 7.4 (low dose) or 18.5 (high dose) kBq of ^225^Ac-NM600, or vehicle IV. Tumor volume, body weight, and survival were monitored three times weekly for 50 days. Humane endpoints were used based on the UW-Madison IACUC guidelines, including tumor volume reaching 700% of the initial volume, significant weight loss (20% of initial body weight), and significant decline in general health.

In addition, in a second round of therapeutic studies, randomized groups of 10 bearing subcutaneous MyC-Cap (FVB/NJ) or Tramp-C1 (C57BL/6) tumors were given 18.5 kBq of ^225^Ac-NM600, 18.5 kBq of ^225^Ac-NM600 + anti-PD1 (200ug, intraperitoneally, three doses three days apart starting 4 days after radiotracer administration), ^225^Ac-NM600 + isotype control (200ug, intraperitoneally, three doses three days apart starting 4 days after radiotracer administration), ^225^Ac-NM600 + anti-CD8 (200ug, intraperitoneally, twice a week throughout the end of the study) or vehicle IV. A schematic representation of therapeutic study groups can be found in **Table S1**.

### Tumor Microenvironment Immunomodulation

#### Flow Cytometry

Tumors from TrampC-1 or MyC-CaP tumor-bearing mice from different treatment groups at different time points were collected (n=3 per group) and immediately processed for flow cytometry analysis. Tumors were chopped and digested at 37° in RPMI1640 mixed with DNAse and protease inhibitor. After Fc blocking (BD, Franklin Lakes, NJ, 553142) for 20 minutes at 4°C, cells were then stained for CD11b, CD25, GR-1, CD3, MHCII, CD45, CD4, CD8, CD44, KLRG-1-PE, CD69, CD62l, CD103, CD27, and Foxp3 according to previously published protocol (*20*). A Thermo Fisher Attune NxT cytometer and FlowJo v10 were used for analysis. Gates were set according to fluorescence minus one (FMO) controls.

#### Immunocytokine Panel

Tumor tissues were collected and flash frozen in liquid nitrogen and then stored at −80°C until background levels were reached. Samples were prepared by chopping into small pieces and lysing with a solution consisting of NP40 detergent supplemented with PMSF and protease inhibitor cocktail for 30 minutes. Samples were centrifuged at 14000 rpm for 10 minutes at 4°C, and the supernatant was extracted for analysis using a ProcartaPlex Cytokine and Chemokine Panel (Waltham, MA) according to manufacturer recommendations. Briefly, a protein standard was reconstituted for serial dilution to create a standard curve with 7 points. Samples were loaded onto a 96-well plate, with bead mix added to all wells, followed by washing. The plate was then covered and shaken for 2 h at 500 rpm. After shaking, the plate was washed, and detection antibody was added prior to shaking for 30min; this step was repeated with Streptavidin-PE. Finally, the reading buffer was added, and the plate was analyzed using a Luminex MAGPIX (Austin, TX). Manufacturer-recommended settings of at least 50 beads collected were used for the analysis.

### Statistical Analysis

All quantitative data were analyzed using GraphPad Prism (version 8.4.2) and are presented as the mean ± standard deviation. Survival curves were displayed using the Kaplan-Meier method and tested for significance using the Log-rank test. Comparisons between groups were made using two-way ANOVA or unpaired *t*-test, where *P* < 0.05 was considered statistically significant.

## RESULTS

### SPECT/CT Imaging reveals elevated and sustained tumor accretion of NM600 in both tumor models

Longitudinal SPECT/CT studies were performed to determine the tumor-targeting and biodistribution of NM600 and to estimate the radiation dosimetry of ^177^Lu-NM600 and ^225^Ac-NM600 in mice bearing Tramp-C1 or MyC-CaP tumors. Longitudinal SPECT/CT images displayed as maximum-intensity projections at different time points can be seen in **Figure 1A**. Quantitative region-of-interest analysis of the SPECT images (**Figure 1B**) revealed initially elevated blood pool activity of the radiotracer 5-h-postinjection and gradual decline with a circulation half-life of 29.6 ± 8.7 h and 17.1 ± 4.1 h for MyC-CaP and Tramp-C1 tumor-bearing mice, respectively (**Figure S1**). Owing to NM600’s hepatobiliary excretion, liver uptake peaked at 24 h post-injection (p.i.) and declined over time. Probe accumulation in off-target tissues, such as kidneys, spleen, bone, and muscle, was below 3 %IA/g at all measured time points. Elevated tumor accumulation was observed at early timepoints, with peak values of 7.9 ± 1.3 and 5.0 ± 1.3%IA/g in Tramp-C1 and MyC-CaP tumors, respectively. Tumor uptake persisted with high uptake in the tumor still seen at 72 h p.i. *Ex vivo* biodistribution studies carried out after the last SPECT time point in **Figure S2** agreed with the SPECT findings, with an uptake of 6.6 ± 1.2 and 7.8 ± 1.3 %IA/g in Tramp-C1 and Myc-CaP tumors, respectively.

**Figure 1.**
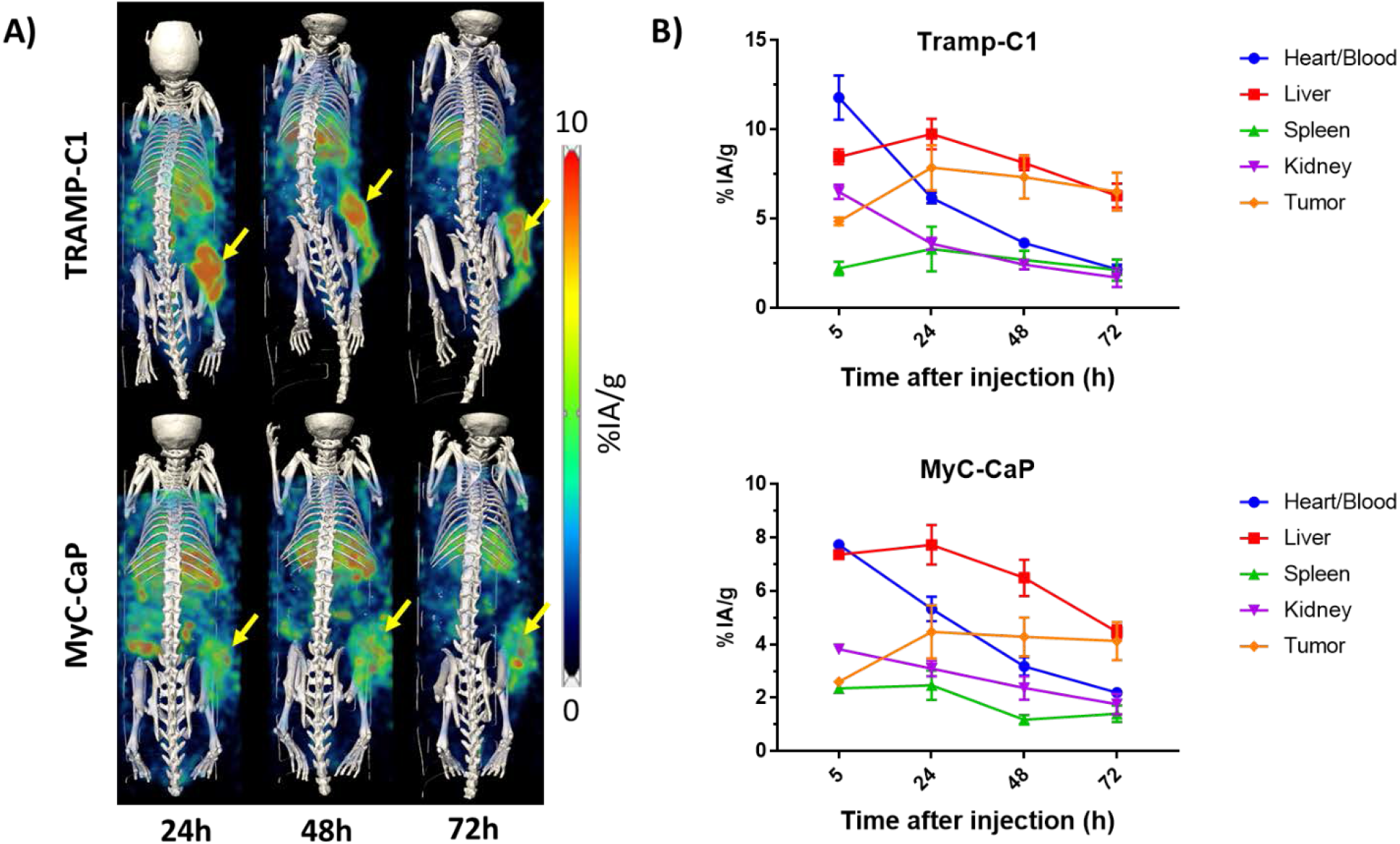
Longitudinal ^177^Lu-NM600 SPECT/CT imaging in murine models of prostate cancer**. A)** Maximum-intensity projections of Tramp-C1 and Myc-CaP tumor-bearing mice at 24, 48, and 72 hours after i.v injection of ^177^Lu-NM600 (n=3) demonstrates clear tumor uptake throughout the study and hepatobiliary excretion of the probe. **B)** Time-activity curves with quantitative region-of-interest analysis of SPECT imaging demonstrate uptake in the tumor reached 7.9 ± 1.3 and 5.0 ± 1.3%IA/g in Tramp-C1 and MyC-CaP tumors, respectively. Data are presented as %IA/g (mean ± SD).

### Serial *ex vivo* biodistribution studies confirm findings from SPECT and demonstrate that choice of isotope does not affect NM600 biodistribution

To further understand the pharmacokinetics and biodistribution dynamics of NM600 radiolabeled with both alpha and betta emitters, biodistribution studies of up to 192h p.i were carried out. 3.7 MBq of ^177^Lu-NM600 (0.27 μg/MBq) or 7.4 KBq of ^225^Ac-NM600 (2.7 μg/MBq) were intravenously administered in Tramp-C1, or MyC-CaP tumor-bearing mice and organs were collected at different time points. Target and off-target tissue uptake were similar in animals injected with ^177^Lu-NM600 and ^225^Ac-NM600, which were also in agreement with findings from SPECT/CT (**Figure S3**). Tumor uptake was highest at 24 h with 10.0 ± 2.5 and 8.3 ± 2.7 %IA/g for ^177^Lu-NM600 and at 72h with 7.4 ± 3.3 and 4.2 ± 0.9 %ID/g for ^225^Ac-NM600 in Tramp-C1 and MyC-CaP tumors respectively. The off-target tissue with the highest uptake was the liver due to hepatobiliary excretion of the probe. ^225^Ac-NM600, but not ^177^Lu-NM600 liver uptake increased over time, consistent with previously published reports for ^225^Ac-DOTA compounds (*27*).

### Dosimetry estimations indicate delivery of comparable or higher radiation doses to the tumor than to all normal tissues

Dosimetry estimates were calculated for ^177^Lu-NM600 based on longitudinal SPECT/CT using our in-house developed Monte Carlo voxel-based dosimetry method (*26*), and data were presented as Gy/MBq of injected activity. Tumors received the highest dose of all tissues in both models, with 1.1 and 0.8 Gy/MBq to Tramp-C1 and MyC-CaP tumors, respectively (**Table S2**). Normal tissue dosimetry showed similar trends, with the liver receiving the highest off-targeted dose among all organs.

Due to the observed discrepancies in liver uptake with ^177^Lu-NM600, ^225^Ac-NM600 dosimetry was estimated based on longitudinal *ex vivo* biodistribution data. The absorbed dose estimates in Gy/kBq for ^225^Ac-NM600 in mice bearing either TRAMP-C1 or MyC-CaP tumors are summarized in **Table S3**. Tumor absorbed doses of 0.58, and 0.25 Gy/kBq were estimated for Tramp-C1 and MyC-CaP tumors, respectively. The liver received the largest dose of all normal organs with 1.32 and 0.90 Gy/kBq in Tramp-C1 and MyC-CaP bearing animals, respectively, which can be attributed to the hepatobiliary clearance of the agent. The remaining normal tissues had markedly lower absorbed dose values well under 0.90 Gy/kBq, including the bone marrow, with 0.10 and 0.07 Gy/kBq, in Tramp-C1 and MyC-CaP tumors, respectively.

### Toxicity studies evidenced ^177^Lu/^225^Ac NM600 tolerability

After identifying absorbed doses in the tumor and off-target organs, we wanted to investigate if the administered doses were safe and if potential toxicities had arisen. Administration of ^177^Lu-NM600 or ^225^Ac-NM600 had a negligible impact on animal weight (**Figure S4**) and survival, suggesting the high tolerability of the treatments. Additionally, we further established safety profiles for ^177^Lu-NM600 and ^225^Ac-NM600 through a panel of cellular, histological, and functional tests evaluating hematological, renal, and hepatic toxicity.

Complete blood counts (CBC) analyses of Tramp-C1 and MyC-CaP tumor-bearing mice administered high or low ^177^Lu-NM600 or ^225^Ac-NM600 injected activity evaluated potential hematologic toxicity (n = 5; **Figure 2**). In all treated groups, animals had transient, dose-dependent reductions in white and red blood cells and lymphocytes, indicative of mild bone marrow toxicity. In addition, animals presented with moderate leukopenia that resolved at later timepoints. Slight anemia was observed at earlier time points but resolved within 2 weeks. Overt thrombocytopenia, lower platelet levels compared to controls, occurred on day 14 for Tramp-C1 and day 28 for MyC-CaP tumor-bearing mice. In general, even at the highest IA level, the effects were well tolerable.

**Figure 2.**
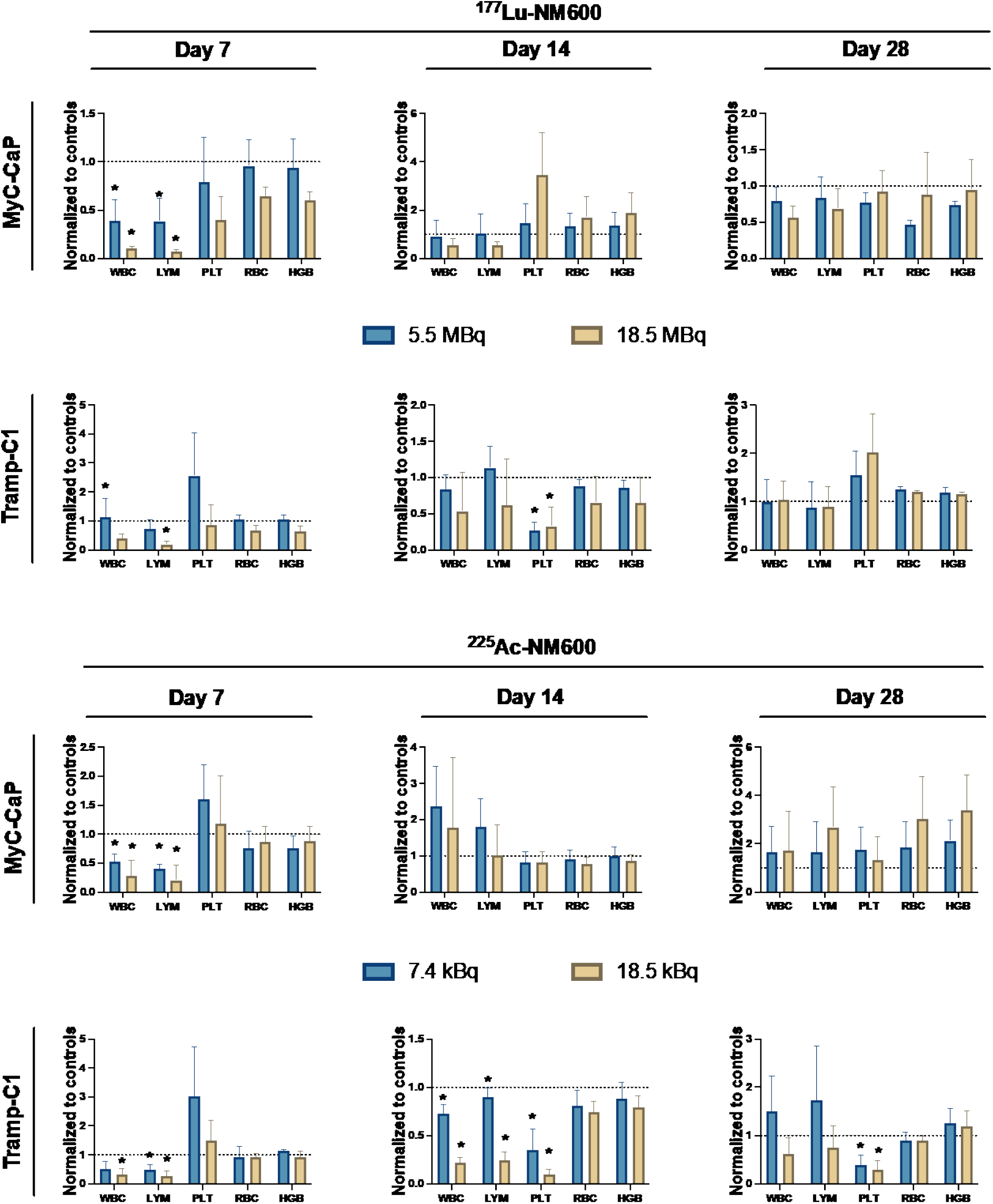
Complete Blood Count panels of MycCaP and TrampC1 tumor-bearing mice (n = 5) on days 7, 14, and 28 after administration of a high or low dose of ^225^Ac-NM600 or ^177^Lu-NM600. Compared to controls, animals that received RPT presented transient, dose-dependent reductions in hematological parameters, including white blood cells, lymphocytes, and red blood cells, indicative of mild bone marrow toxicity. Values returned to baseline by day 28 and did not result in animal lethality. Values are normalized to controls. *p < 0.05.

A comprehensive metabolic panel analysis of 15 parameters indicative of acute kidney and liver toxicity (**Figures S5**) was carried out on days 7, 14, and 28 after RPT administration (n=3). Normal levels of liver and kidney function readouts were observed in MyC-CaP and TrampC-1 tumor-bearing animals following ^177^Lu-NM600 (**Figure S5A**) and ^225^Ac-NM600 treatments (**Figure S5B**). Interestingly, mice treated with ^177^Lu-NM600 displayed lower AST values than controls on day 28. MyC-CaP tumor-bearing mice had significantly (p<0.05) lower BUN levels at day 28 after administration of a low or high dose of ^225^Ac-NM600. Tramp-C1 tumor-bearing exhibited significantly (p<0.05) lower ALT levels at both ^225^Ac-NM600 injected activity levels. Even though significantly lower ALT, AST, and BUN levels were found at later time points, these values do not usually correlate with kidney or liver toxicity. However, mice treated with a high IA also presented significantly (p<0.05) lower globulin levels at day 28, which can indicate kidney or toxicity.

To uncover potential toxicity in radiation-sensitive organs, we performed histological (H&E) examinations of several organs (**Figures S6-S9)**. The liver, kidneys, and spleen showed no signs of overt tissue degeneration in any of the time points investigated (days 7, 14, or 28) in either tumor model, doses or radioisotopes administered. Liver and kidney sections also presented normal morphology. H&E staining of the bone marrow revealed reduced cellularity for both doses of ^225^Ac-NM600 or ^177^Lu-NM600 in both tumor models on day 7, but normal morphology on day 28. Interestingly, intestines of Tramp-C1 and MyC-CaP tumor-bearing mice receiving a high dose of 177Lu-NM600 showed signs of inflammation with the presence of immune infiltrates present at earlier timepoints (days 7 and 14) but more prominent at later time points (day 28). Immune infiltrates were also present in the intestines of MyC-CaP tumor-bearing mice that received 18.5kBq ^225^Ac-NM600. Intestines of animals administered 5.55 MBq ^177^Lu-NM600 or 7.4 kBq ^225^Ac-NM600 presented normal morphology and no signs of inflammation. Overall, these results warrant further treatment optimization and the exploration of potential long-term toxicities.

### ^225^Ac-NM600, but not ^177^Lu-NM600, promotes anti-tumor effects and increases overall survival in both tumor models

In therapeutic studies using ^177^Lu-NM600, animals received a single 5.55 MBq (low) or 18.5 MBq (high) IV injection delivering 6.1 or 20.5 Gy to Tramp-C1, and 4.5 or 15.3 Gy to MyC-CaP tumors, respectively. Similarly, 7.4 kBq (low) and 18.5 kBq (high) ^225^Ac-NM600 delivered 4.3 and 10.7 Gy to Tramp-C1 and 1.8 or 4.6 Gy to MyC-CaP tumors, respectively. Treatment with ^177^Lu-NM600 resulted in only modest dose-dependent inhibition in TRAMP-C1 tumor growth (**Figure 3A**) and no significant survival advantage compared to controls (**Figure 3B**). Myc-CaP-bearing mice were also resistant to ^177^Lu-NM600 treatment with no detectable differences in tumor growth inhibition and survival between groups (**Figure 3A-B**). In contrast, significant (p < 0.01) tumor growth inhibition was observed at either administered ^225^Ac-NM600 activity level compared to control in both MyC-Cap and Tramp-C1 tumors (**Figure 3A**). Tumor doubling times for control and ^225^Ac-NM600, 7.4 kBq or 18.5 kBq treatment groups were 4.2, 7.2, and 15.4 in MyC-CaP bearing animals, and 6.0, 13.1, and 14.8 days for animals bearing Tramp-C1 tumor, respectively. Notably, a significant (p < 0.0001) extension in overall survival (**Figure 3B**) was achieved for both tumor models in animals treated with ^225^Ac-NM600 7.4 kBq or 18.5 kBq. Median survival of MyC-CaP mice was extended from 12 d in control animals to 43 and 51 d in the ^225^Ac-NM600 7.4 kBq or 18.5 kBq groups, respectively, while Tramp-C1 mice showed median survivals of 27 and 63 d for control and 7.4 ^225^Ac-NM600; median survival was not reached in the ^225^Ac-NM600 18.5 kBq treatment group.

**Figure 3.**
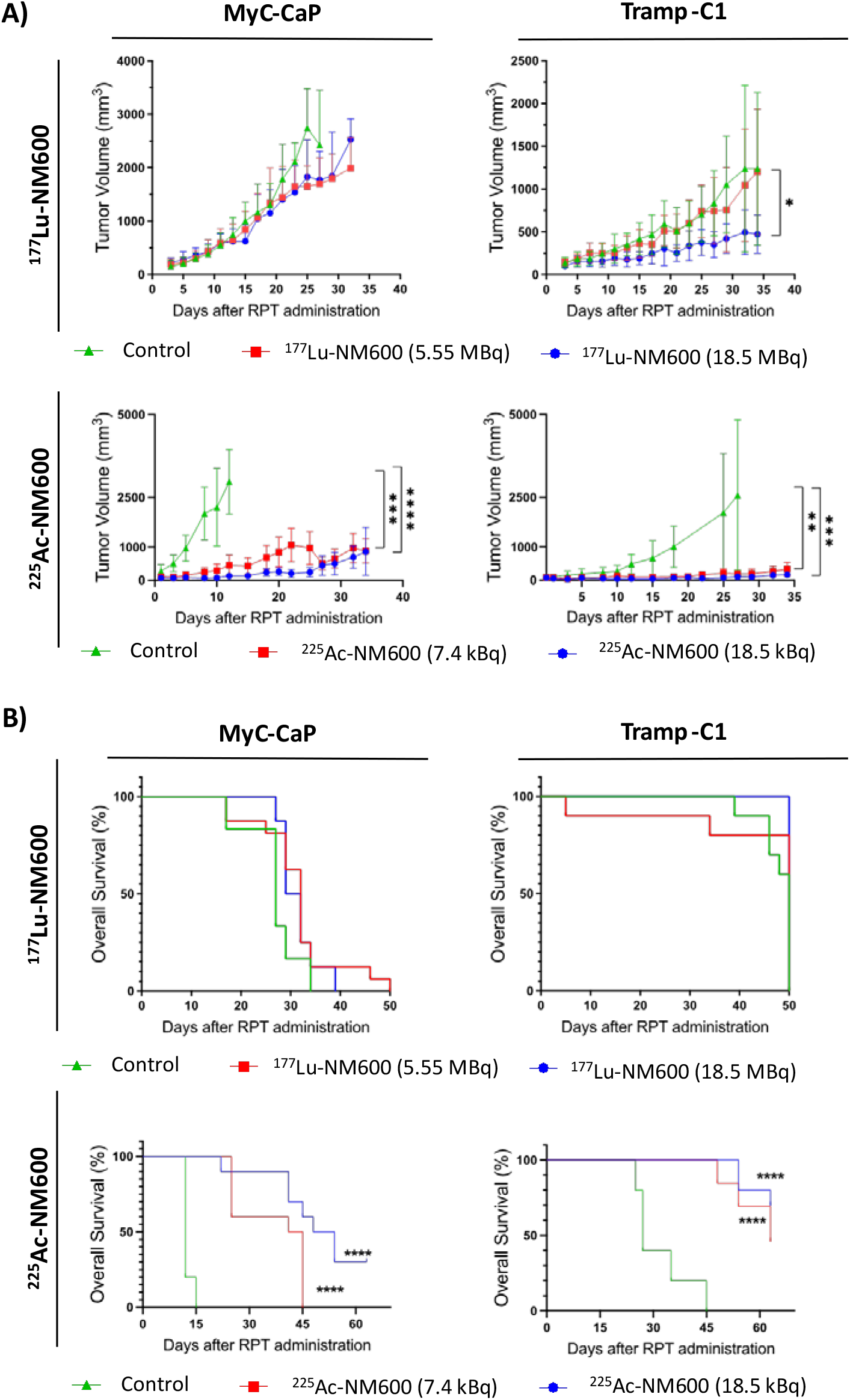
Therapeutic studies (n = 10) of Tramp-C1 and MyC-CaP tumor-bearing mice that received vehicle (control), 5.55 or 18.5 MBq of ^177^Lu-NM600, and 7.4 or 18.5 kBq of ^225^Ac-NM600. **A)** Tumor growth curves demonstrate that treatment with ^177^Lu-NM600 resulted in modest dose-dependent inhibition in Tramp-C1 but no inhibition in MyC-CaP. In contrast, compared to controls, significant tumor growth inhibition was observed at either administered dose of 225Ac-NM600 in both MyC-CaP and Tramp-C1 tumors. **B)** Kaplan-Meier curve demonstrates no improvement in overall survival in either Tramp-C1 or MyC-CaP tumor-bearing animals receiving ^177^Lu-NM600 but significantly improved overall survival in both tumor models given low or high ^225^Ac-NM600 IA. *p < 0.05, **p < 0.01, ***p < 0.001, ****p < 0.0001.

#### TME immune profiling demonstrates that ^225^Ac-NM600 promotes abrogation of infiltrating suppressor cells, a more active CD8 repertoire, and activation of effector and memory T cells

We next performed detailed immunophenotyping of suppressive and effector cells within TRAMP-C1 and MyC-Cap tumor microenvironment. Treatment with ^177^Lu-NM600 resulted in a significant increase in infiltrating suppressive immune cells, including myeloid-derived suppressive cells (MDSC: CD45+CD11b+GR-1+) (p = 0.0003) for the high dose group at day 28 when compared to control) and regulatory T cells (Tregs: CD4+CD25+FoxP3+) (p = 0.0005 for high dose group at day 28 when compared to control) in TrampC-1 tumors (**Figure 4 A, B, G**) and MDSCs in MyC-CaP tumors (p = 0.0008 at day 28 when compared to controls) (**Figure 4 D**). Interestingly, in TrampC1 tumors, 18.5 MBq of ^177^Lu-NM600 significantly increased both MDSCs (p = 0.0206 at day 28) and Tregs (p = 0.0297 at day 28) compared to 5.55MBq.

**Figure 4.**
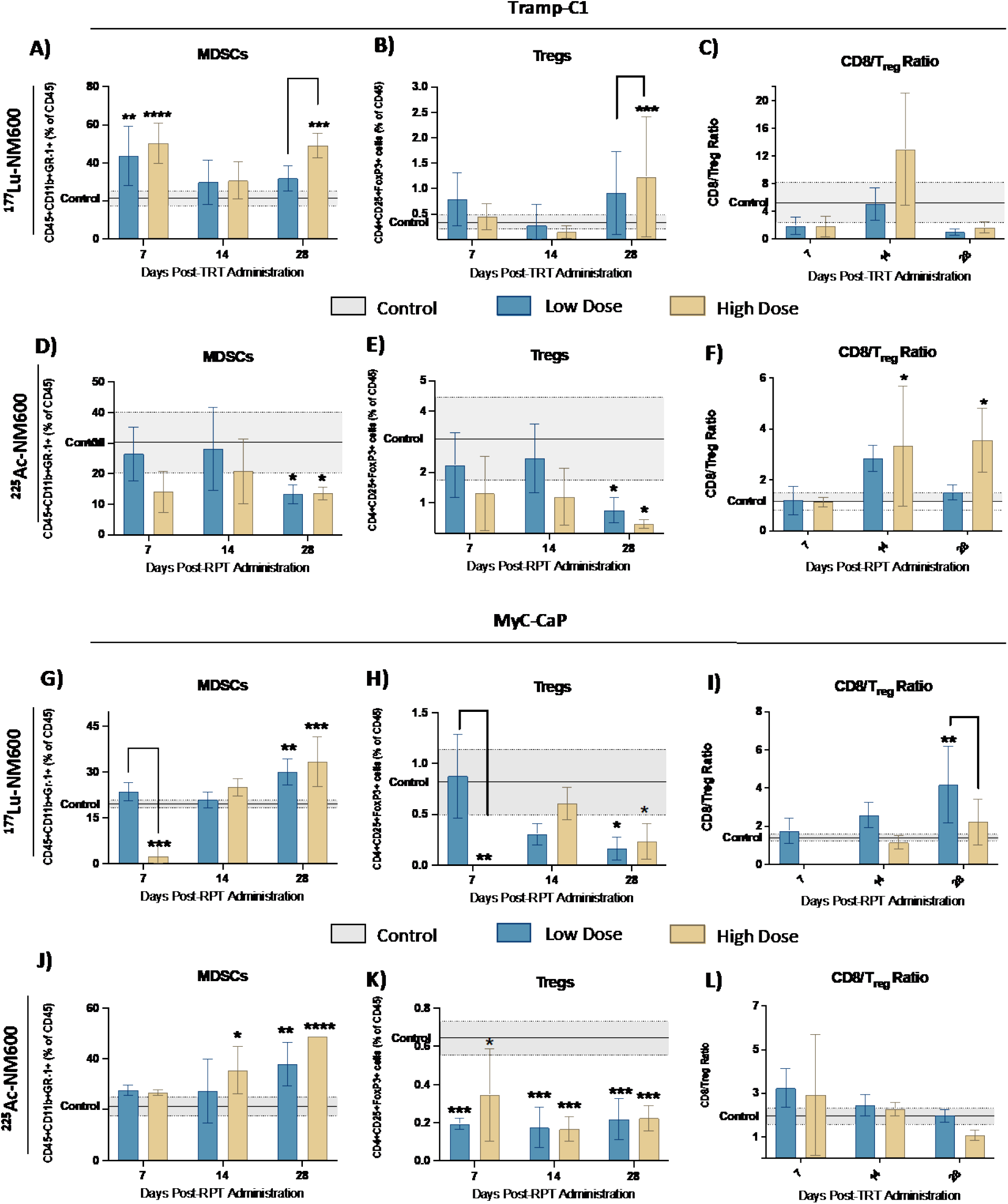
^225^Ac-NM600 decreases levels of immunosuppressive cells such as MDSCs and Tregs while promoting a higher CD8/Treg ratio. Flow Cytometry analysis of (A, D) Myeloid-derived suppressor cells (MDSCs), (B, E) Regulatory T Cells (Tregs), and (C, F) CD8/Treg ratio of Tramp-C1 tumors injected with low (5.5 MBq) or high dose (18.5 MBq) of ^177^Lu-NM600 or low (7.4 kBq) or high dose (18.5 kBq) of ^225^Ac-NM600. (G, J) Myeloid-derived suppressor cells (MDSCs), (H, K) Regulatory T Cells (Tregs), and (I, L) CD8/Treg ratio of MyC-CaP tumors injected with low (5.5 MBq) or high dose (18.5 MBq) of ^177^Lu-NM600 or low (7.4 kBq) or high dose (18.5 kBq) of ^225^Ac-NM600. Statistical analysis when compared to controls or otherwise noted. * p< 0.05, ** p<0.01, *** p<0.001.

The ratio of cytotoxic T cells to Tregs (CD8/Treg ratios) showed a dynamic behavior over the 4-week study period but ultimately led to decreased CD8/Treg ratios at day 28 when compared to controls (**Figure 4C**) in TrampC1 tumor-bearing animals injected with ^177^Lu-NM600. On the contrary, ^225^Ac-NM600 administration resulted in a significant reduction (p<0.05) in both MDSC and Tregs, leading to a steady increase in CD8/Treg ratio (p = 0.0171 at day 28) in animals receiving 18.5 kBq ^225^Ac-NM600 (**Figure 4F**). Significant reduction in infiltrating Tregs was also observed with 7.4 kBq ^225^Ac-NM600 but only on day 28 post-treatment. Treg drove overall changes in CD8/Treg ratios as CD8+ T cells levels in TrampC1 tumor were largely unaltered by both ^177^Lu-NM600 or ^225^Ac-NM600 treatment at all investigated time points (**Figure S10 C and F)**. RPT significantly decreased CD3 and CD4+ T cells in both tumor models, but that effect was more pronounced in MyC-CaP animals (p < 0.0001) that received 18.5MBq of ^177^Lu-NM600, probably because of the high absorbed dose (15Gy) **(Figure S10 G, H, J, K)**.

We next wanted to investigate the effects of RPT on cytotoxic T cells. In both tumor models, CD44+ memory cells, activation (CD69 expression), and the number of proliferating cells (Ki67+) markers remained mostly unaltered with the administration of ^177^Lu-NM600 (**Figure 5 A-C and G-I**). In contrast, administration of ^225^Ac-NM600 promoted a more active CD8 repertoire with significantly increased levels of CD69 (p = 0.0042) at day 28, 18.5kBq) and Ki67 (p = 0.0193) at day 28, 18.5kBq) in TrampC1 (**Figure 5 E, F**) and CD69 (p < 0.0001 at day 28, 7.4 kBq), Ki67 (p = 0.0235) at day 28, 18.5kBq) and CD44 (p = 0.0152 at day 28, 18.5kBq) in Myc-CaP tumors (**Figure 5 J-L).** We also investigated CD8 memory activity (**Figure S11**) and found that effector memory (CD27-CD62L-), central memory (CD27+CD62L+), and resident memory (CD69+CD103+) remained mostly unaltered after RPT administration in Myc-CaP. Markers on CD8+ cells were significantly increased, especially at earlier timepoints, in TrampC-1 tumors following ^225^Ac-NM600 but unaltered after ^177^Lu-NM600 (**Figure S11 A-N**). In addition, short-lived effector cells (KLRG-1CD127+) were unaffected by ^225^Ac-NM600 but were significantly increased in Tramp-C1 tumor-bearing mice while significantly decreased (p < 0.0001) in MyC-CaP tumors by ^177^Lu-NM600 (**Figure S11 D, H, L, P**).

**Figure 5.**
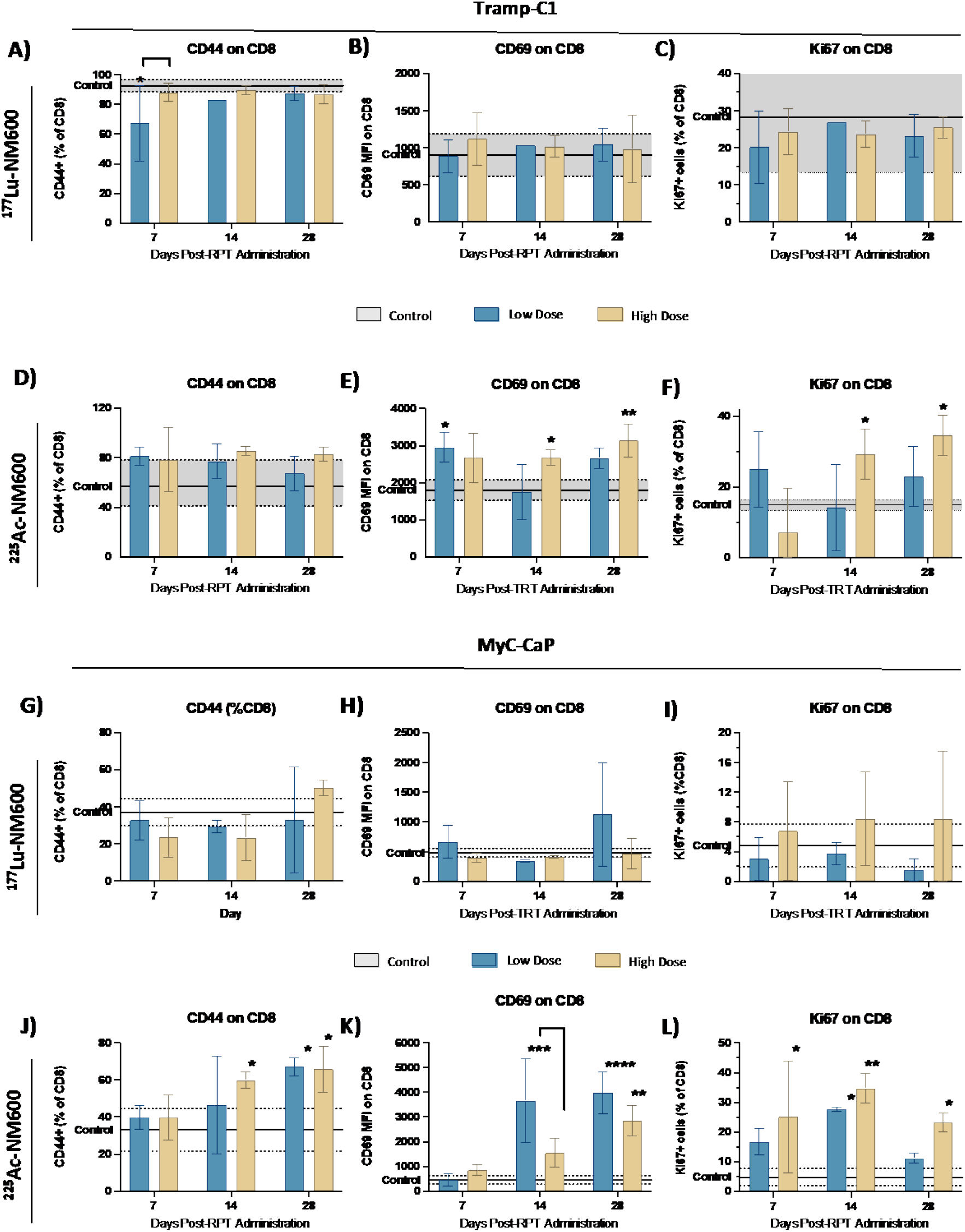
^225^Ac-NM600 promotes a more activated CD8+ repertoire with increased levels of activation/memory (CD44+), activation (CD69+) and proliferation (Ki67+) markers. Flow cytometry analysis in Tramp-C1 tumor-bearing mice of **(A)** CD44+, (B) CD69+, and (C) Ki67+ after administration of ^177^Lu-NM600 and (**D)** CD44+, **(E)** CD69+, and **(F)** Ki67+ after administration of ^225^Ac-NM600. Flow Cytometry analysis in MyC-CaP tumor-bearing mice of **(G)** CD44+, **(H)** CD69+, and **(I)** Ki67+ after administration of ^177^Lu-NM600 and **(J)** CD44+, **(K)** CD69+, and **(L)** Ki67+ after administration of ^225^Ac-NM600. Data are presented as % of CD8+ T cells. Statistical analysis when compared to controls or otherwise noted. * p< 0.05, ** p<0.01

#### ^225^Ac-NM600 augments PD-1 expression on CD8+ T cells and PD-L1 expression on myeloid cells

In Tramp-C1 tumors, no statistical difference in PD-1 levels on CD8+ T cells was found between control and groups injected with either ^177^Lu-NM600 injected dose (**Figure 6A**). In contrast, significantly (p<0.0001) increased PD-1 expression on CD8+ T cells was found in TrampC-1 tumors of animals injected with 18.5 kBq ^225^Ac-NM600 on day 14 (34.8 ± 2.8 % of CD8) and 7.4 kBq ^225^Ac-NM600 on day 28 (32.1 ± 5.8 % of CD8) compared to controls (7.5 ± 4.7 % of CD8+) (**Figure 6B**). In MyC-CaP tumor-bearing mice, 18.5kBq ^225^Ac-NM600 and 18.5MBq ^177^Lu-NM600 significantly (p<0.05) elevated PD-1 expression on CD8+ T cells were also found on day 14 when compared to controls (**Figure 6 C,D**). We also found that PD-L1 expression on MDSCs (**Figure 6 E-H**) significantly increased with RPT, with maximum values 2-fold higher than those found for controls in MyC-CaP tumor-bearing mice injected with ^225^Ac-NM600 (**Figure 6H**).

**Figure 6.**
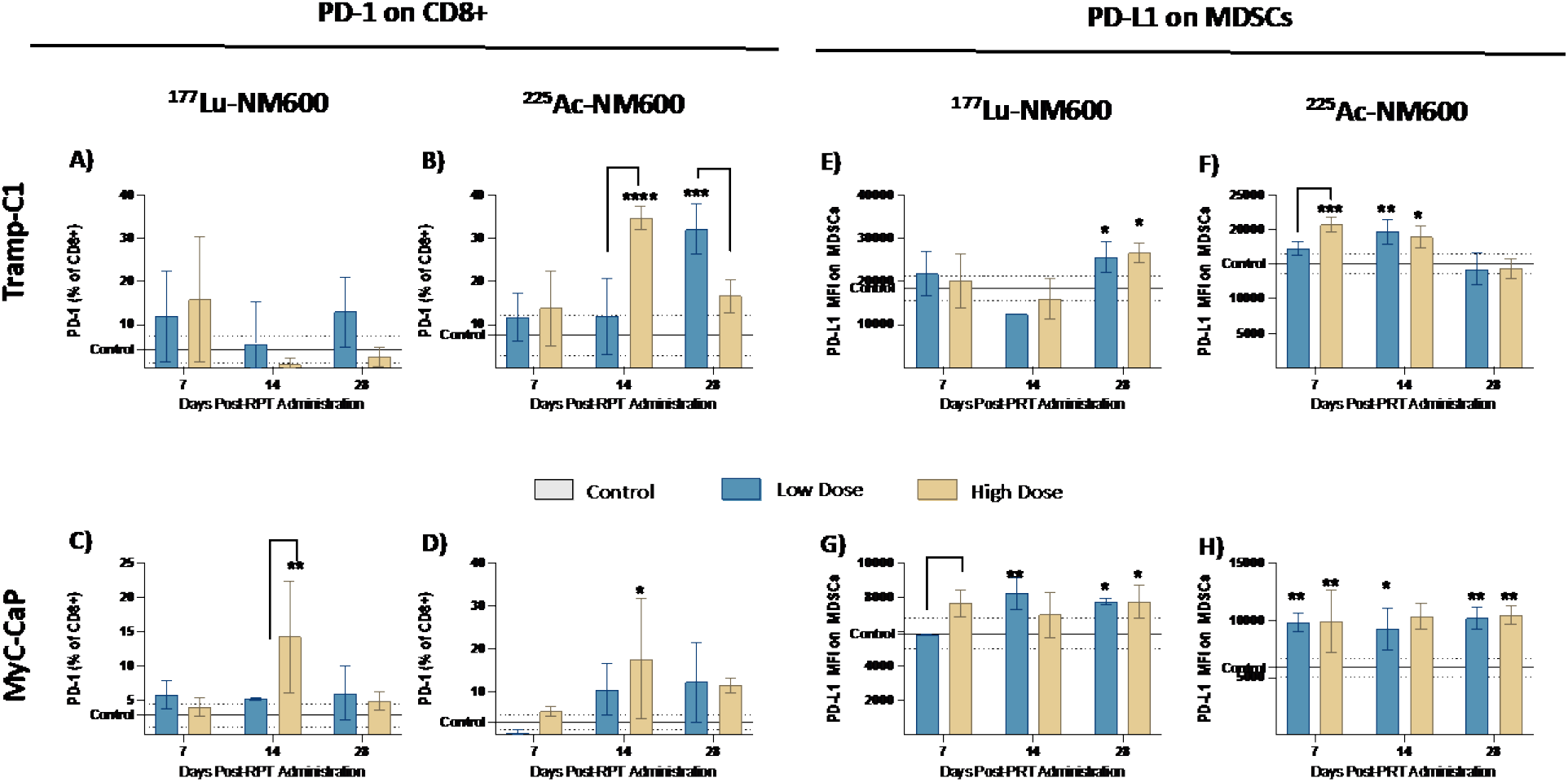
Flow Cytometry results indicate that ^225^Ac-NM600 increases PD-1 expression on CD8+ T cells and PD-L1 expression on MDSCs. PD-1 expression on CD8+ cells after administration of **(A)** ^177^Lu-NM600 or **(B)** ^225^Ac-NM600 in Tramp-1 and of **(C)** ^177^Lu-NM600 and **(D)** or ^225^Ac-NM600 in MyC-CaP tumor-bearing mice. PD-L1 expression on MDSCs in animals injected with **(E)** ^177^Lu-NM600 and **(F)** or ^225^Ac-NM600 in Tramp-C1 tumor-bearing mice and with **(G)** ^177^Lu-NM600 and **(H)** or ^225^Ac-NM600 in MyC-CaP tumor-bearing mice. Statistical analysis when compared to controls or otherwise noted. *p<0.05, ** p<0.01, *** p<0.001, ****p<0.0001

Since the combination of RPT and PD-1/PD-L1 inhibitors has been extensively proposed, we sought to investigate if there were radiation-induced PD-L1 expression changes in tumor cells (**Figure S12**). ^225^Ac-NM600 administration did not affect PD-L1 expression of Tramp-C1 or MyC-CaP tumors; however, administration of ^177^Lu-NM600 promoted a marked decrease (p<0.01) in PD-L1 expression in Tramp-C1 tumors when compared to controls (Figure **S16A**).

#### Cytokine analyses reveal that higher activity levels increase chemokines and pro-inflammatory cytokines and that ^225^Ac-NM600 also decreases suppressive cytokines

To uncover potential differences in cytokine profiles between ^177^Lu-NM600 and ^225^Ac-NM600, we analyzed 26 cytokines and chemokines in tumor tissue collected at different time points (days 7, 14, and 28) post-RPT administration. **Figure 7** shows a heatmap of our findings, plotted as fold change from controls. Overall, lower ^177^Lu-NM600 or ^225^Ac-NM600 injected activities reduced cytokine and chemokine levels in Tramp-C1 tumors, while cytokine and chemokine levels were increased at a higher injected activity. Fold changes were accentuated with ^225^Ac-NM600 compared to ^177^Lu-NM600. In MyC-CaP tumors, we uncovered intriguing differences between the isotopes investigated, albeit less pronounced. For example, IL-4 and IL-23 levels increased with ^177^Lu-NM600 but decreased with ^225^Ac-NM600. In addition, concentrations of most chemokines were much higher in the ^225^Ac-NM600 high dose group than in the other groups investigated. To look for overall trends in cytokine and chemokine levels across different doses and isotopes, we organized analytes by function into Th1, Th2, Th17, and chemokines.

**Figure 7.**
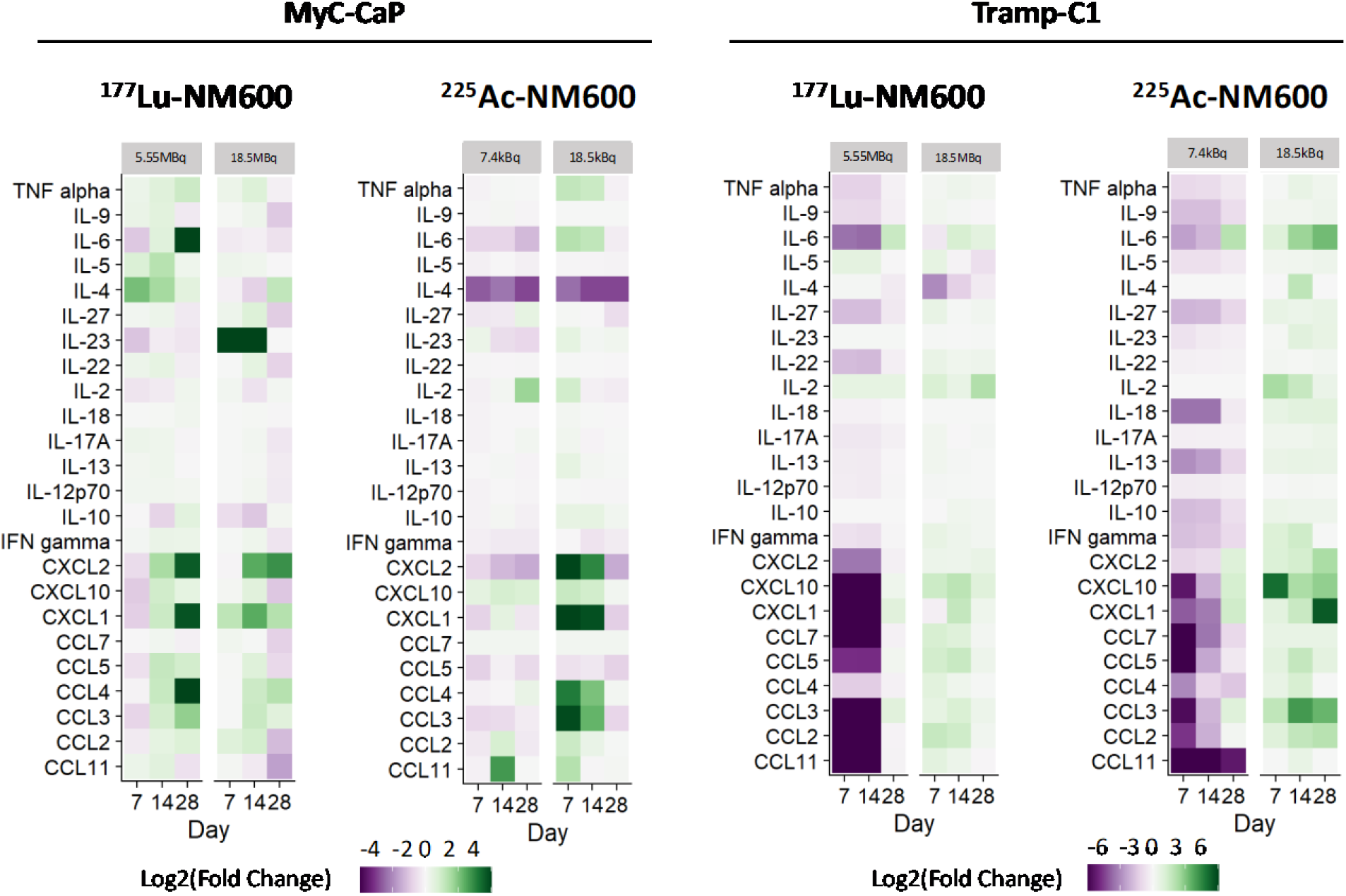
Heatmap representing cytokine and chemokine concentration changes in MyC-CaP and Tramp-C1 tumors were analyzed on days 7, 14, and 28 after RPT administration. Data is presented as Log2(Fold Change) compared to controls (n = 3). In Tramp-C1 tumors, ^225^Ac-NM600 (7.4kBq) or ^177^Lu-NM600 (5.55MBq) decreased cytokine and chemokine levels while ^225^Ac-NM600 (18.5kBq) or ^177^Lu-NM600 (18.5MBq) increased their levels. In MyC-CaP tumors, some cytokines such as IL-4 and IL-23 increased with ^177^Lu-NM600 but decreased with ^225^Ac-NM600, while most chemokines significantly increased with an 18.5 kBq of ^225^Ac-NM600. Concentration changes were more prominent with ^225^Ac-NM600 than ^177^Lu-NM600.

##### Th1 phenotype

Overall, protein targets of the Th1 phenotype remained mostly unaltered in MyC-CaP tumors but had dose-dependent variations in TrampC1 tumors, with a high dose of any of the isotopes increasing its levels while lower doses promoted decreased levels. In MyC-CaP tumors, levels of IFN gamma, IL-12p70, IL-18, TNF, IL-27, and IL-2 remained mostly unaltered after RPT (data not shown). In TrampC1 tumors (**Figure S17**), a low dose of either ^225^Ac-NM600 or ^177^Lu-NM600 significantly decreased IFN gamma, IL-12p70, IL-27, and TNF alpha values while a high IA of either ^225^Ac-NM600 or ^177^Lu-NM600 significantly increased them. IL-2 levels on trampC1 tumors significantly increased in animals that received a higher dose of ^225^Ac-NM600 when compared to controls on days 7 and 14, but no statistical difference was found between any other groups at any of the time points investigated.

##### Th2phenotype

Similarly to Th1 phenotype changes, proteins of the Th2 phenotype changed more in TrampC1 tumors than in MyC-CaP tumors. Concentration changes were dose-dependent, with a lower dose of both isotopes promoting decreased levels, while a high dose promoted slightly higher or unaltered concentration levels (**Figure S14**). ^225^Ac-NM600, at a lower dose, was more efficient in decreasing Th2 proteins than ^177^Lu-NM600. For example, IL-9 levels significantly decreased (reaching a p < 0.0001) in animals administered with 7.4 kBq ^225^Ac-NM600 and 5.55MBq ^177^Lu-NM600 but significantly increased (p < 0.05) in animals injected with 18.5 kBq ^225^Ac-NM600 or 18.5MBq ^177^Lu-NM600 (**Figure S14D**). We observed the same trend for IL-10 and IL-13 (**Figure S14 E-F**), with the administration of 18.5 kBq ^225^Ac-NM600 promoting increased levels while 7.4kBq ^225^Ac-NM600 promoted decreased levels.

##### Th17pphenotypes

IL-17A and IL-22 followed a similar trend as other cytokines, and in TrampC1 tumors (**Figure S19A-B**), a lower dose of ^177^Lu-NM600 or ^225^Ac-NM600 promoted diminished levels when compared to controls and significantly increased levels when compared to a high dose of either isotope. While ^177^Lu-NM600 did not promote any changes in IL-23, 7.5kBq ^225^Ac-NM600 decreased its levels, and 18.5kBq increased its levels when compared to the baseline control. In contrast to Th1 and Th2 cytokines, ^177^Lu-NM600, but not ^225^Ac-NM600, seemed to alter Th17 cytokines in MyC-CaP tumor-bearing mice at later time points, promoting significantly (p < 0.05) diminished levels of IL-17A and IL-22 (**Figure S15 D, E**). In addition, 18.5 MBq of ^177^Lu-NM600 increased (p < 0.01) IL-23 levels when compared to all other groups investigated (**Figure S15F**).

##### Chemokines

We observed similar trends in chemokine changes (low dose RPT diminishes while high dose RPT augments) (**Figure S20**), with 5.55 MBq ^177^Lu-NM600 or 7.4kBq ^225^Ac-NM600, yielding significantly reduced levels of CXCL10, CXCL2, CXCL1, CCL2, CCL7, CCL3, CCL4, CCL5, and CCL11 when compared to all other groups.

#### ^225^Ac-NM600 RPT combined with anti-CD8 or anti-PD1 treatments does not affect anti-tumor response

Given the immunomodulatory effects of ^225^Ac-NM600 in both Tramp-C1 and MyC-Cap, we investigated the impact of CD8+ T cell depletion and anti-PD1 checkpoint inhibitions on the anti-tumor efficacy of high IA of ^225^Ac-NM600. Administration of 18.5 kBq ^225^Ac-NM600 and a CD8+ cell-depleting antibody twice weekly throughout the study did not abrogate ^225^Ac-NM600 anti-tumor efficacy (**Figure S17**). However, no statistically significant difference in tumor growth inhibition between mice given ^225^Ac-NM600, ^225^Ac-NM600 + anti-CD8, or ^225^Ac-NM600 + isotype-matched control. Similarly, combination studies administering 18.5 kBq ^225^Ac-NM600 and anti-PD-1 antibody on days 4, 7, and 10 post-RPT administration had negligible effects on ^225^Ac-NM600 tumor growth inhibition (Figure S21). Overall, these results further suggest that ^225^Ac-NM600 efficacy is less dependent on the infiltration or stimulation of cytotoxic CD8+ T cells than on the reduction of suppressive lineages within the TME.

## DISCUSSION

Most studies have focused on maximizing the radiation dose delivered to prostate tumor cells using a maximum tolerable dosing approach based on normal tissue toxicity. However, this paradigm assumes that a tumor cannot be overdosed, generally ignoring the presence and importance of the tumor microenvironment. Overwhelming preclinical and clinical data indicate that anti-tumor effects of radiation are partly mediated by radiation-induced immunological effects (*28*). Radiation therapy (RT) can enhance immune susceptibility by promoting cell death that leads to tumor-neoantigen presentation and ultimately T cell activation, releasing of chemokines that promote T-cell infiltration, and temporarily depleting of RT-sensitive immune lineages, including suppressor and effector T-lymphocytes (*25, 29–32*). These effects depend on RT dose and fractionation and manifest at tumor-absorbed doses as low as 4 Gy of low LET radiation (*24, 33*). It has been shown that proper radiation dose must be delivered to all tumor sites (*23*) since untreated tumors can affect systemic and local immune responses and hamper anti-tumor effects (*34*), a phenomenon called concomitant immune tolerance (CIT). While simultaneously delivering absorbed doses at primary and distant tumor sites is unattainable with conventional radiotherapy (EBRT), RPT can potentially overcome CIT by irradiating the collective tumor burden. Unfortunately, most of our understanding of RPT radiobiology derives from preclinical studies in immunocompromised animals, which lack the immunological dimension of an anti-tumor response. Furthermore, radiopharmaceutical therapy studies have rarely employed a dosimetry-guided theranostic approach; therefore, knowledge gaps exist surrounding the absorbed dose and radioisotope dependencies of RPT immunomodulation and toxicity.

Since the effects of alpha-emitting RPT agents in the tumor microenvironment of prostate cancer are relatively unknown, we aimed to investigate not only the potential of NM600 as a targeted agent for prostate cancer but also how different isotopes (^177^Lu and ^225^Ac) and absorbed doses can affect tumor immunology and drive anti-tumor response in two different immunocompetent prostate cancer models. NM600’s unique ability to target lipid rafts, which are overexpressed by nearly all cancer cells, including murine prostate cancers, and be radiolabeled with a variety of imaging and therapeutic radioisotopes enabled us to perform studies impractical with other RPT agents. The *theranostic* capacity of NM600 to be used for *therapeutic* delivery of RT and *diagnostic* imaging was critical to studying the radiobiological effects of RPT. To this end, we leveraged the gamma emission of ^177^Lu to perform *in vivo* SPECT-CT imaging, enabling monitoring of the tumor-selective uptake and retention of ^177^Lu-NM600. We then employed the image-derived biodistribution data to perform voxel-based Monte Carlo dose estimations to the tumor and pertinent normal tissues (e.g., spleen and bone marrow) using an in-house platform (RAPID) (*26, 35*). This permitted absorbed dose-based comparisons between the different agents and dosing regimens, essential to dissecting the mechanistic underpinnings of RPT immunomodulation.

SPECT/CT imaging and biodistribution studies show semi-selective tumor targeting and prolonged retention for ^177^Lu-NM600 and ^225^Ac-NM600 in both MyC-CaP and Tramp-C1 tumor models. Treatment with up to 18.5 MBq (500 μCi) ^177^Lu-NM600 did not lead to significant anti-tumor effects or survival benefits in any of the tumor models, evidencing the relative radioresistance of these tumors to ^177^Lu β-radiation. In contrast, both administered activity levels of ^225^Ac-NM600 significantly (p < 0.001) inhibited tumor growth and extended mean overall survival (p < 0.0001) by at least 3-fold in both animal models. In addition, anti-tumor effects were more pronounced in TrampC-1 tumors than MyC-CaPs mice, in line with dosimetry findings showing a 2x higher absorbed dose per unit of IA for Tramp-C1 versus MyC-Cap tumors. More interestingly, the lower ^225^Ac-NM600 IA (7.4 kBq) delivering 4.3 and 1.8 Gy to Tramp-C1 and MyC-CaP tumors, respectively, was more efficacious than high ^177^Lu-NM600 IA imparting 20.5 and 15.3 Gy to these tumors, respectively. These results indicate relative biological effectiveness (RBE) for ^225^Ac-NM600 exceeding 4.8 and 8.5 in Tramp-C1 and MyC-CaP tumors, which are markedly higher than the reported RBE range of 2-5 for ^213^Bi and ^211^At α-radiation (*36*). An important caveat to these studies was the use of murine cancer models in an immunodeficient host, which excluded the critical contribution of the immune system to tumor control. Although establishing revised RBE values for ^225^Ac in the context of a fully functional immune system is outside of the scope of the present work, our findings showed significant and distinctive radiation quality-dependent immune effects that warrant such endeavors in the near future.

The normal tissue biodistribution of ^177^Lu-NM600 and ^225^Ac-NM600 led to significant absorbed doses to the liver. Although absorbed doses to the liver were kept below 25 Gy, well under the reported 65 Gy mean dose to the whole liver for patients with no severe complications following ^90^Y-microsphere radioembolization (*37*). We also carried out a series of radiotoxicity evaluations to elucidate the toxicity profile. Serial histological examination of potential organs at risk of radiotoxicity, such as the heart, liver, spleen, and kidney, revealed no signs of acute cellular toxicity. In addition, via CBC analysis, we identified mild bone marrow toxicity with dose-dependent cytopenia on early time points which were confirmed through histology of bone marrow; however, these were resolved by day 28, similar to our previous findings (*20–22*). Functional kidneys and liver metabolic assays further confirmed ^177^Lu-NM600 and ^225^Ac-NM600’s tolerability, with no clinically relevant effects on blood chemistries at any of the IA or time points investigated. Overall, treatments were tolerated, and the bone marrow appears to be dose-limiting. Interestingly, histology revealed the presence of immune infiltration in the intestine of animals that received higher IA of either ^225^Ac-NM600 or ^177^Lu-NM600. Plausibly, these could result from intestinal irradiation, given the hepatobiliary excretion of the agent resulting in radiation-induced intestinal inflammation, a typical response following abdominal irradiation in the clinic (*38*). Furthermore, systemic immune activation can lead to adverse events such as colitis which also presents as intestinal immune infiltration (*39–41*). Further investigation on the extra tumoral immune effects of RPT are necessary to determine potentially deleterious effects, especially at high IA levels, on the systemic immune system that may undermine mounting of an anti-tumor immune response or potential synergies with immunotherapies.

Absorbed tumor doses alone could not explain the large differences in tumor control and survival between ^177^Lu-NM600 and ^225^Ac-NM600 treatments. We found that immunomodulatory effects of RPT caused by different isotopes could potentially be partly responsible for different treatment outcomes *in vivo*. Immune system responses directly related to cells and secreted mediators in the prostate cancer TME can dictate treatment outcomes (*42, 43*). Among infiltrating cells, CD8+ T cells are ultimately responsible for tumor cell lysis through granules and cytokines release. Even though a higher level of cytotoxic T cells usually indicates greater anti-tumor effects (*44–46*), that is not necessarily true for prostate cancer (*47*). Some studies have reported that an increase in cytotoxic T lymphocytes (CTL) in prostate cancer TME correlated with a worse prognosis (*48–50*), mainly because CTLs were neutralized by anti-inflammatory and immunosuppressive mediators such as MDSCs and Tregs (*50, 51*). Thus, generally, a metric more reflective of the immunological status of the TME is the ratio of infiltrating CTLs to suppressive cells (*52*). In fact, decreased CD8/Treg ratio is generally recognized as a marker of poor outcomes in solid tumors (*53–55*). In this study, we observed a similar phenomenon where ^177^Lu-NM600 treatment led to higher CD8+ TCL infiltration but worse tumor control than ^225^Ac-NM600. This seemingly contradictory result is satisfactorily explained by the significant increase in MDSCs and Tregs after administration of ^177^Lu-NM600, which resulted in reduced CD8+/Tregs and CD8+/MDSC ratios compared to baseline. These results suggest an inability of β-radiation to eradicate regulatory T cells and other immunosuppressive cell lineages, which are relatively radioresistant compared to their CTL counterparts (*31, 56*). We have recently shown similar results using another β-emitter, where the accumulation of Tregs within the TME negated the anti-tumor response to ^90^Y-NM600 treatment and led to antagonistic effects in combination with anti-PD1 immunotherapy (*20*). Tumor control was then rescued through Treg depletion using an anti-CTLA4 antibody, evidencing the significant impact of Tregs on the tumor radiobiology of these prostate tumors.

On the contrary, despite ^225^Ac-NM600 treatment resulting in a slightly reduced CD8+ infiltrate, the profound depleting effects on suppressive lineages driven by α-particles led to overall pro-inflammatory immune cell balance and enhanced anti-tumor efficacy; CD8/Treg ratios were significantly higher after ^225^Ac-NM600 when compared to ^177^Lu-NM600 and controls. These findings point to previously unbeknownst advantages of ^225^Ac high-LET radiation to abrogate immunosuppressive populations in prostate cancer. Other cellular effects, including CTL senescence and anergy also contribute to the relatively immunologically cold phenotype of the prostate cancer TME (*57–60*). Notably, we observed that ^225^Ac-NM600 but not ^177^Lu-NM600 induced a more active T cell repertoire, with increased expression of activation and proliferation markers such as CD44+, CD69+, and Ki67+ on CD8+ T cells, which contributed to generating an immunologic “hot” TME. Since ^225^Ac-NM600 anti-tumor efficacy seemed driven by predominantly different immunomodulatory effects compared to ^177^Lu-NM600: increased CD8+/Treg ratio in Tramp-C1 tumor-bearing mice or more active CD8+ T cells repertoire in MyC-CaP, we wanted to investigate whether effector CD8+ T cells mediated anti-tumor efficacy. Concomitant ^225^Ac-NM600 and CD8 depletion showed negligible influence on the anti-tumor response of ^225^Ac-NM600, suggesting that Treg/MDSC depletion rather than CD8+ activity was the main driver of improved tumor control efficacy.

Besides cellular components, mediators such as cytokines and chemokines play a prominent role in pro- or anti-tumor properties (*61*). Under proper stimulation, CD4+ T cells differentiate into Th1, Th2, and Th17 phenotypes. Th1 cells produce pro-inflammatory cytokines such as IFN-γ and IL-12, which exert anticancer activity and are linked to antigen presentation and tumor cytotoxicity (*62*). In contrast to the Th1 response, Th2 cells mediate humoral immunity and produce IL-4, IL-10, and other cytokines that can promote cancer progression and have been linked to angiogenesis, immune suppression, and tumor proliferation (*63*). The role of the third subset of helper T cells, Th17, is still controversial since they are characterized by the production of pro-inflammatory cytokines such as IL-17A and IL-22 but have also been shown to promote tumor growth in prostate cancer (*64–66*). Employing a Luminex-based assay, we investigated the TME cytokine profile of prostate cancer models following RPT. In Tramp-C1 tumors, Th1, Th2, Th17, and chemokines all collectively decreased with administration of low IA but increased at high IA of either ^177^Lu-NM600 or ^225^Ac-NM600. Interestingly, a low IA of RPT seemed to decrease both pro-inflammatory and suppressive cytokines/chemokines in Tramp-C1 tumors, while high RPT IAs seemed to increase both. Since high RPT IA appeared to exert a better anti-tumor response than low IA, most likely, an increase in pro-inflammatory cytokines had a more pronounced effect than an increase in suppressive mediators, although this effect can be confounded by increased direct cell killing at higher administered activities. Further studies are needed to investigate the timing and effects of individual cytokine/chemokine in prostate cancer. In MyC-CaP tumors, changes in cytokines and chemokines were less apparent compared to controls, yet a few interesting changes in mediators were observed. For example, IL-4, which is usually linked to poor prognosis, activation of the androgen receptor, and increased proliferation in prostate cancer (*67*), increased with both high and low ^177^Lu-NM600 IA but decreased with either ^225^Ac-NM600 IA. The same pattern was observed with IL-23, which is secreted by myeloid cells and drives androgen resistance by activating the androgen receptor pathway and promoting cell survival and proliferation in androgen-deficient environments (*68*). Furthermore, chemokines such as CCL3 and CCL4 that drive immune activation were markedly higher in animals receiving ^225^Ac-NM600 compared to ^177^Lu-NM600 in both tumor models. Upon interaction with dendritic cells, CD4+ cells produce CCL3 and CCL4, which are necessary for CD8+T cell activation, initiating a specific immune response (*69, 70*). Increased levels of those chemokines found in ^225^Ac-NM600 tumors suggest immune activation. Overall, compared to ^177^Lu-NM600, ^225^Ac-NM600 had a more robust pro-inflammatory immunomodulatory activity leading to elevated CTL activation markers, reduced regulatory cell populations (Tregs and MDSCs), and increased Th1-biased cytokine signatures, which explain its significantly better anti-tumor response.

Prostate cancer shows all the typical hallmarks of a “cold” tumor, including low mutational burden, minimal T-cell and robust MDSC infiltration, and an overall immunosuppressive tumor microenvironment (TME), contributing to disappointing results in patients with advanced prostate cancer in several clinical trials (*71, 72*). Given the amply documented immunomodulatory effects of EBRT, hundreds of clinical trials are exploring the potential synergistic combination of immunotherapy and radiation therapy. In contrast, because the radio-immunobiology of radionuclides remains largely underexplored, much less is known about the combination of RPT and immunotherapies, especially in the clinical setting. Due to the increased PD-1 levels on CD8+ T cells and PD-L1 levels on MDSCs following ^225^Ac-NM600 RPT, we sought to exploit a potential synergy with anti-PD1 immunotherapy to improve anti-tumor efficacy further, even though anti-PD1 effects have been difficult to be reproduced pre-clinically (*73*). Surprisingly, ^225^Ac-NM600 combined with anti-PD1 had no effects on anti-tumor responses compared to ^225^Ac-NM600 + isotype control or ^225^Ac-NM600 alone, most likely because CD8+ cells were also depleted in that group. These results are in accordance with our recently published studies demonstrating that ^90^Y-NM600 RPT with anti-PD-1 ICI had no benefits over RPT alone in these models unless Tregs were simultaneously depleted (*20*). While these preliminary results showed that, under the tested conditions, the addition of anti-PD1 to ^225^Ac-NM600 did not enhance therapeutic effects, further optimization of this combined regimen may uncover synergies between the two therapeutic modalities. To this end, future studies will investigate the impact of different absorbed dose levels, alternative timing of therapeutic administration, other RPT agents (e.g., ^225^Ac-PSMA-617), and blockades of other checkpoint molecules. Nonetheless, this report highlights that the immunological effects of radionuclides are non-trivial and that combinational therapies employing RPT and ICI need thorough investigation in relevant immunocompetent mouse models to unearth both potential synergies or counterproductive immunomodulatory effects.

## Acknowledgment

This work was partly supported by the Department of Defense (Early Investigator Award, W81XWH1910285) and the NIH National Cancer Institute (PO1CA250972). The content is solely the responsibility of the authors and do not necessarily represent the official views of the NIH or DoD. The authors would like to acknowledge the Cancer Center Support Grant: NCI P30 CA014520, the University of Wisconsin Small Animal Imaging & Radiotherapy Facility, and NIH S10ID028670-01 for supporting this work. The isotopes used in this research were supplied by the U.S. Department of Energy Isotope Program, managed by the Office of Isotope R&D and Production.

