## Supplementary Information for "Profound immunomodulatory effects of ^225^Ac-NM600 drive enhanced anti-tumor response in prostate cancer"

Address: 1111 Highland Ave, 7137, Madison, WI 53705

**Table S1.** Groups and Doses for Therapeutic Study

| Group | Treatment | Administration |
|---|---|---|
| Control | Vehicle (100µL) | i.v. |
| $^{177}$Lu-NM600 (high dose) | $^{177}$Lu-NM600 (18.5 MBq) | i.v. |
| $^{177}$Lu-NM600 (low dose) | $^{177}$Lu-NM600 (5.5 MBq) | i.v. |
| $^{225}$Ac-NM600 (high dose) | $^{225}$Ac-NM600 (18.5 KBq) | i.v. |
| $^{225}$Ac-NM600 (low dose) | $^{225}$Ac-NM600 (7.4 KBq) | i.v. |
| $^{225}$Ac-NM600 + anti-PD1 | $^{225}$Ac-NM600 (18.5 KBq)<br>Anti-PD1 (200µg) | i.v. day 0<br>i.p. days 4, 7 and 10 |
| $^{225}$Ac-NM600 + anti-CD8 | $^{225}$Ac-NM600 (18.5 KBq)<br>Anti-CD8 (200µg) | i.v. day 0<br>i.p. twice a week |
| $^{225}$Ac-NM600 + isotype | $^{225}$Ac-NM600 (18.5 KBq)<br>IgG (200µg) | i.v. day 0<br>i.p. days 4, 7 and 10 |

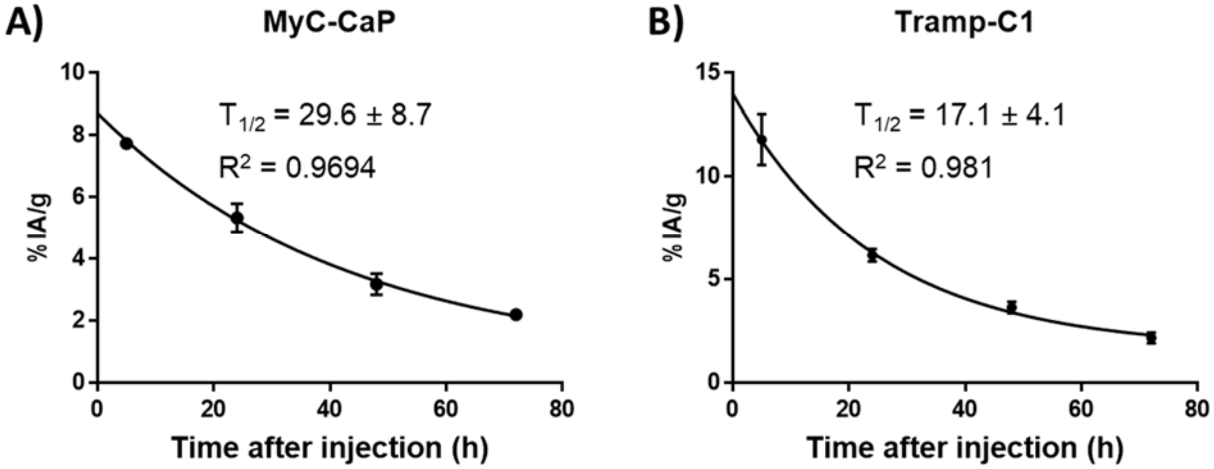


**Figure S1.** Blood time-activity curves of $^{177}$Lu-NM600 in TrampC1 and Myc-CaP tumor-bearing mice. Blood circulation half-life ($T_{1/2}$) was calculated by fitting the quantitative SPECT data to a monoexponential decay. Similar circulation $T_{1/2}$ were observed (29.6 ± 8.7 h vs. 17.1 ± 4.1 h) between the two groups (n=4).

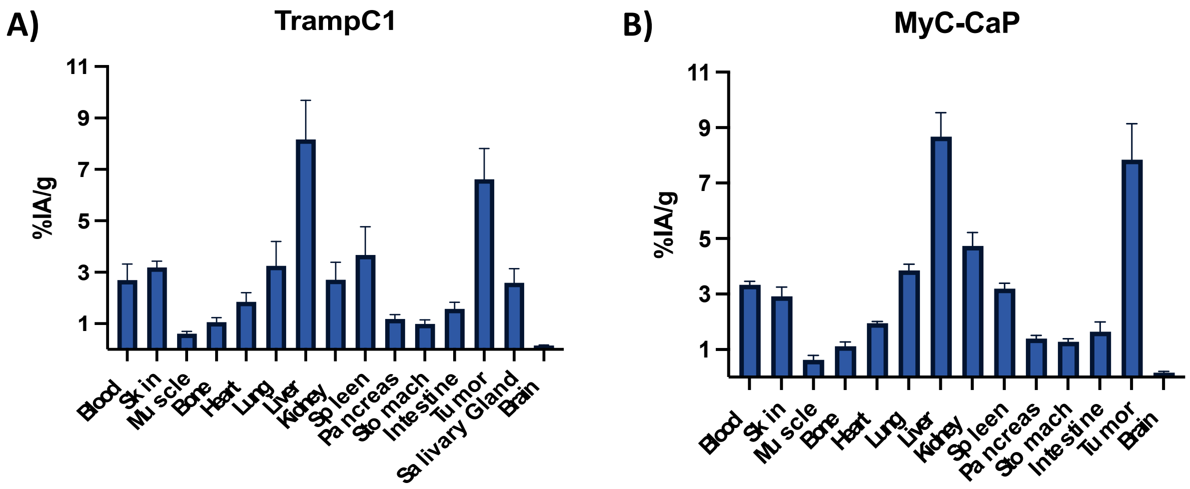


**Figure S2.** Ex vivo Biodistribution studies (n=4) carried out after the last SPECT/CT scan (66h after radiotracer administration) agreed with findings from SPECT in A) TrampC1 and B) MyC-CaP tumor-bearing mice. Data are presented as %IA/g (mean ± SD).

**Table S2**. Summary of results from the dosimetry estimation for $^{177}Lu$-NM600 using longitudinal $^{177}Lu$-NM600 SPECT/CT data and The Standard Mouse Model.

| $^{177}Lu$-NM600 | Tramp-C1 | | | Myc-CaP | | |
|---|---|---|---|---|---|---|
| **Target Organ** | **Absorbed Dose (Gy/MBq)** | **Abosrbed dose (Gy) for low dose** | **Abosrbed dose (Gy) for high dose** | **Absorbed Dose (Gy/kBq)** | **Abosrbed dose (Gy) for low dose** | **Abosrbed dose (Gy) for high dose** |
| Left Kidney | 0.38 | 2.09 | 7.03 | 0.41 | 2.255 | 7.585 |
| Right Kidney | 0.435 | 2.3925 | 8.0475 | 0.476667 | 2.621667 | 8.818333 |
| Liver | 1.2075 | 6.64125 | 22.33875 | 0.976667 | 5.371667 | 18.06833 |
| Spleen | 0.4025 | 2.21375 | 7.44625 | 0.31 | 1.705 | 5.735 |
| Heart | 0.54 | 2.97 | 9.99 | 0.543333 | 2.988333 | 10.05167 |
| **Tumor** | **1.1075** | **6.09125** | **20.48875** | **0.826667** | **4.546667** | **15.29333** |

**Table S3**. Summary of results from the dosimetry estimation for $^{225}$Ac-NM600 using ex vivo Biodistribution data.

| $^{225}$Ac-NM600 | Tramp-C1 | | | Myc-CaP | | |
|---|---|---|---|---|---|---|
| **Target Organ** | **Absorbed Dose (Gy/kBq)** | **Abosrbed dose (Gy) for low dose** | **Abosrbed dose (Gy) for high dose** | **Absorbed Dose (Gy/kBq)** | **Abosrbed dose (Gy) for low dose** | **Abosrbed dose (Gy) for high dose** |
| Bone | 0.016497 | 0.122075 | 0.305187 | 0.105174 | 0.778286 | 1.945716 |
| Heart | 0.16147 | 1.194881 | 2.987202 | 0.122706 | 0.908024 | 2.270059 |
| Lung | 0.748686 | 5.540275 | 13.85069 | 0.488808 | 3.617178 | 9.042944 |
| Liver | 1.32227 | 9.784795 | 24.46199 | 0.902563 | 6.678963 | 16.69741 |
| Kidney | 0.34495 | 2.552627 | 6.381567 | 0.229162 | 1.695798 | 4.239494 |
| Spleen | 0.885477 | 6.552533 | 16.38133 | 0.371722 | 2.750746 | 6.876865 |
| Pancreas | 0.133421 | 0.987313 | 2.468282 | 0.097037 | 0.718074 | 1.795185 |
| Stomach | 0.662689 | 4.903898 | 12.25975 | 0.460338 | 3.406504 | 8.51626 |
| Intestine | 0.078436 | 0.580427 | 1.451067 | 0.045982 | 0.340268 | 0.850671 |
| **Tumor** | **0.577983** | **4.277072** | **10.69268** | **0.246437** | **1.823636** | **4.559089** |
| Marrow | 0.103996 | 0.76957 | 1.923925 | 0.075808 | 0.560979 | 1.402447 |

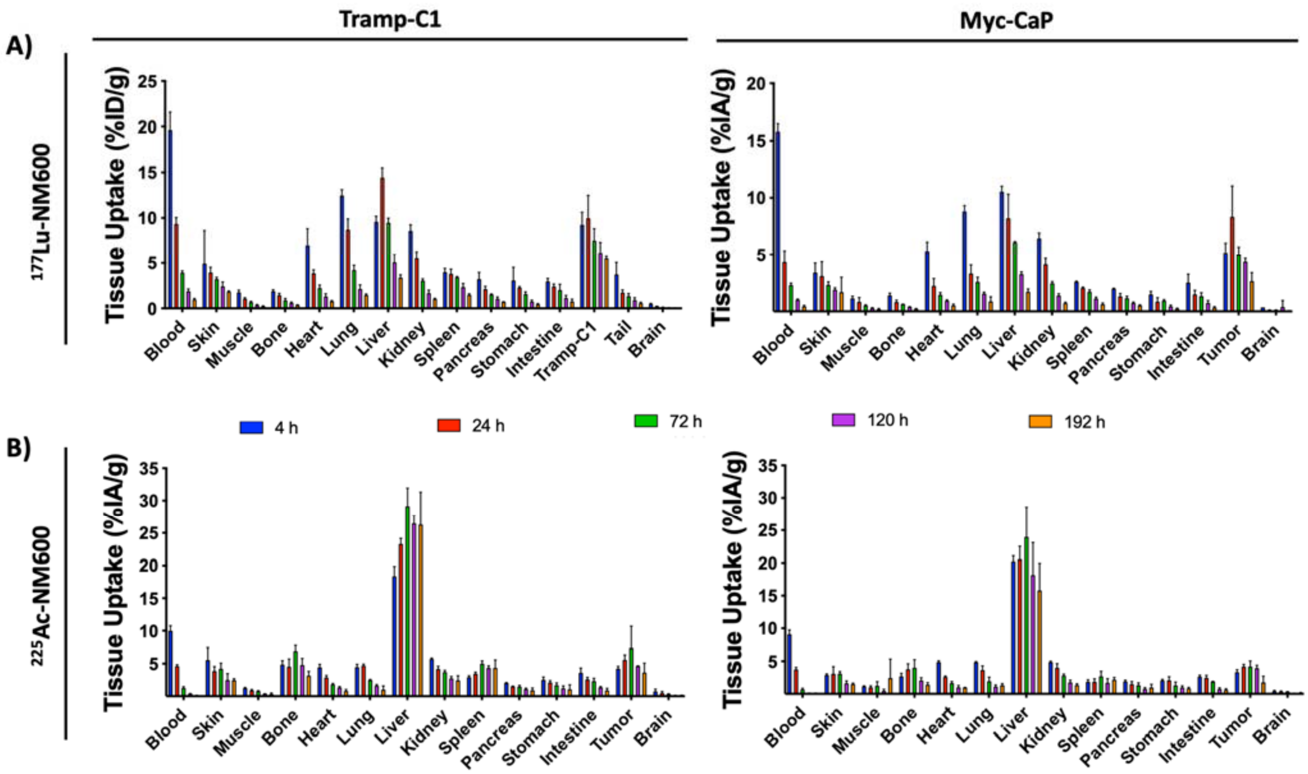


**Figure S3**. Serial *Ex vivo* Biodistribution studies (n=3) in TrampC1 and MyC-CaP tumor-bearing mice after i.v. administration of A) 3.7MBq of $^{177}$Lu-NM600 or B) 7.4 KBq of $^{225}$Ac-NM600. Agreement in tumor uptake and normal-tissue biodistribution was observed between $^{177}$Lu-NM600 and $^{225}$Ac-NM600 injected at the same mass dose, which confirmed the elevated tumor accumulation and retention and the hepatobiliary clearance observed via SPECT/CT imaging. Data are presented as %IA/g (mean ± SD).

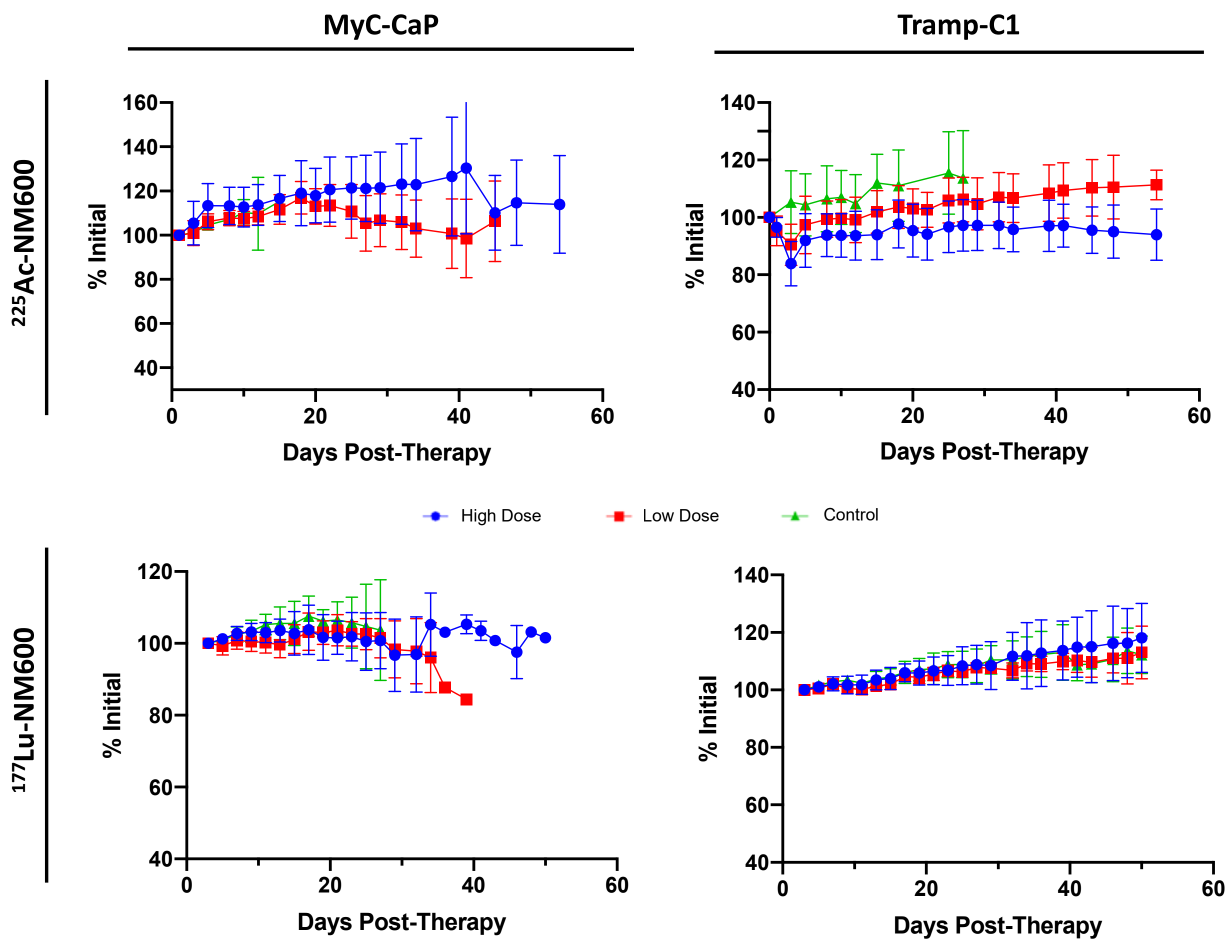


**Figure S4**. Body Weight measurements, represented as percentage of initial body weight of MyC-CaP and Tramp-C1 tumor-bearing mice administered with vehicle or high or low dose of 177Lu-NM600 or $^{225}$Ac-NM600. No significant differences were found between controls and any of the other groups investigated, suggesting high tolerability of the treatments.

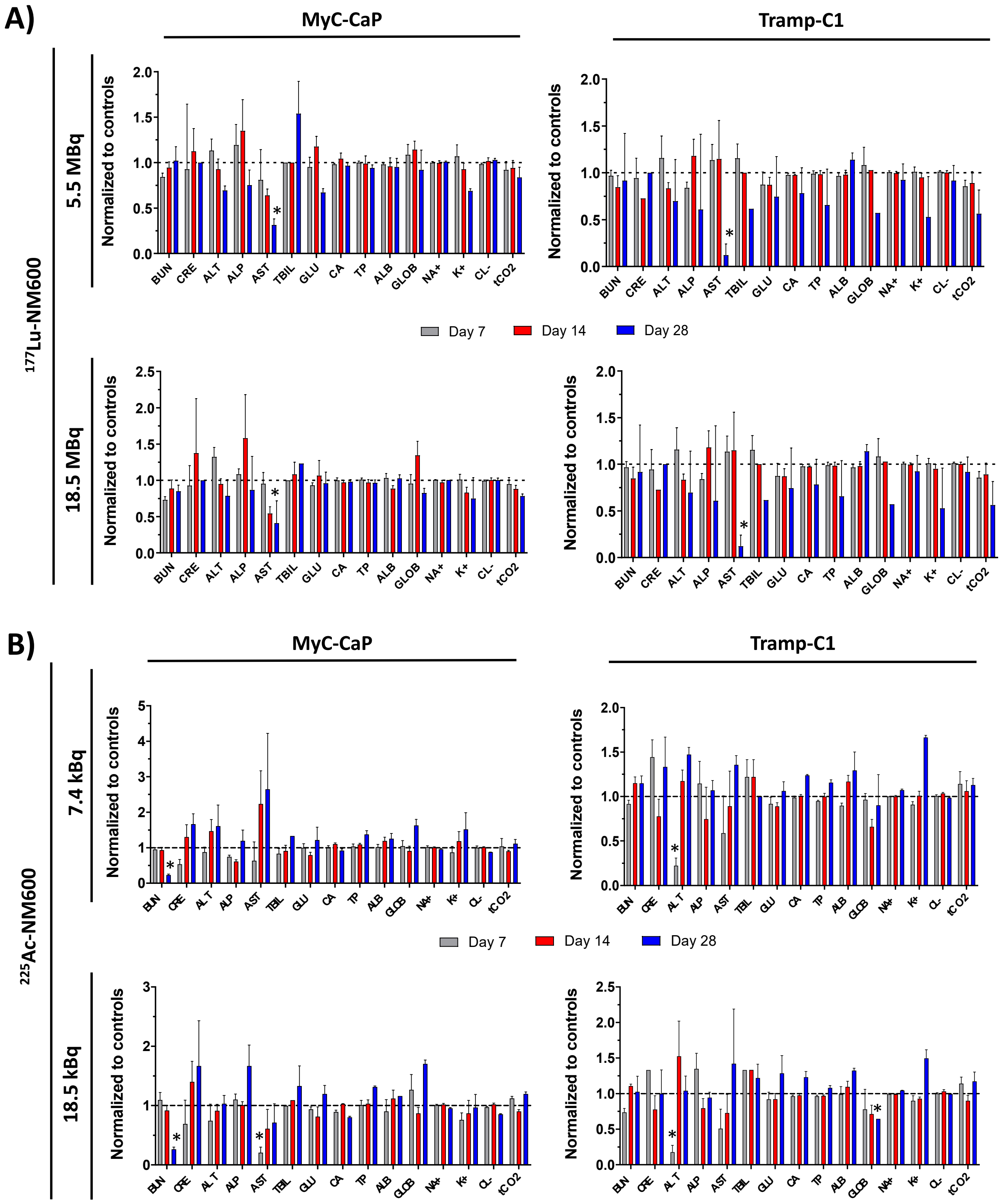


**Figure S5**. Comprehensive Metabolic Panel of MycCaP and TrampC1 tumor-bearing mice at days 7, 14, and 28 after administration of 5.55 MBq or 18.5 MBq of $^{177}$Lu-NM600 or 7.4 kBq or 18.5 kBq of $^{225}$Ac-NM600. Values are normalized to controls. $*p < 0.05$.

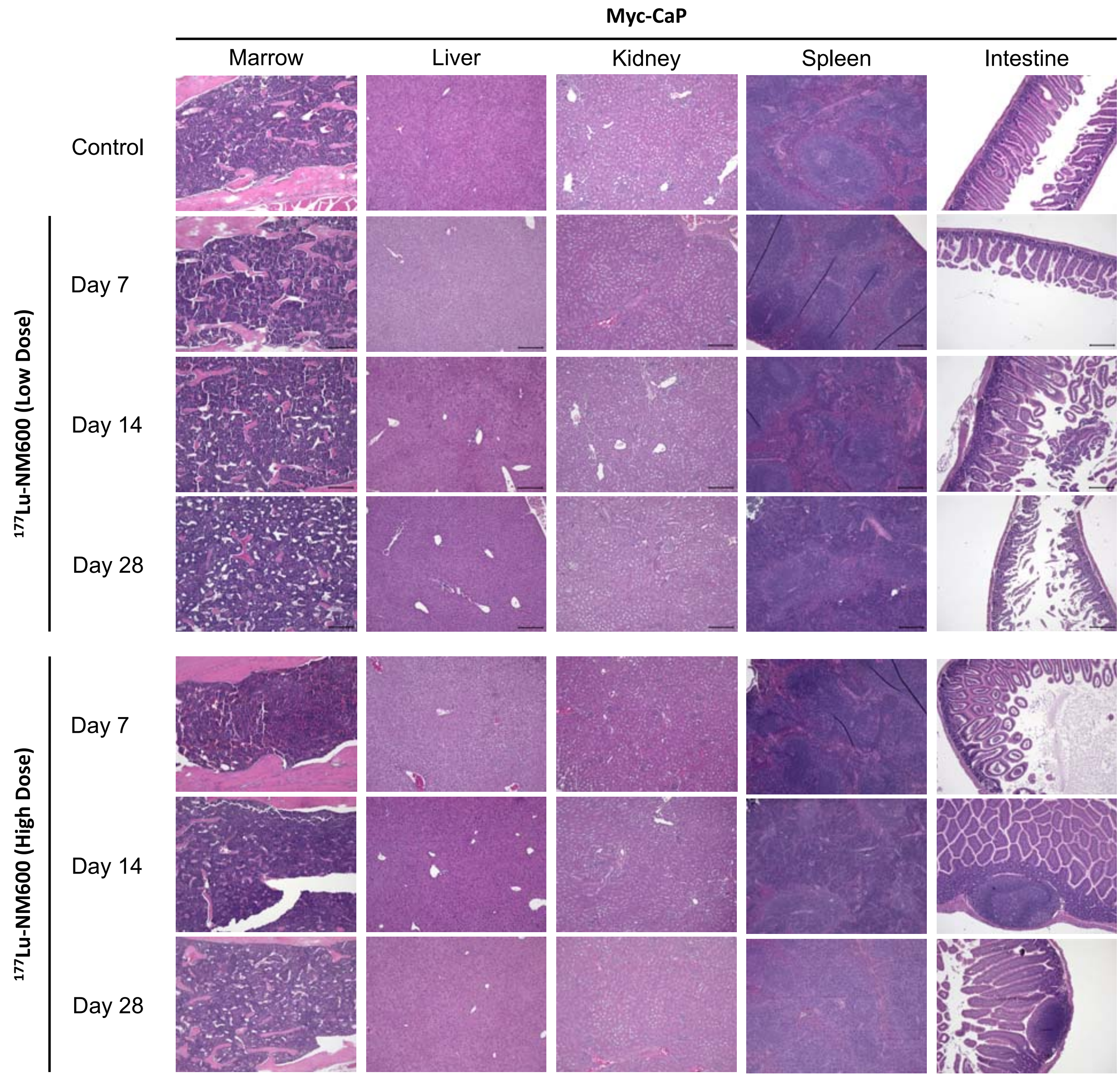


**Figure S6.** H&E staining of major organs collected 7, 14, or 28 days after administration of 5.55 MBq or 18.5 MBq of $^{177}$Lu-NM600 in Myc-CaP tumor-bearing mice.

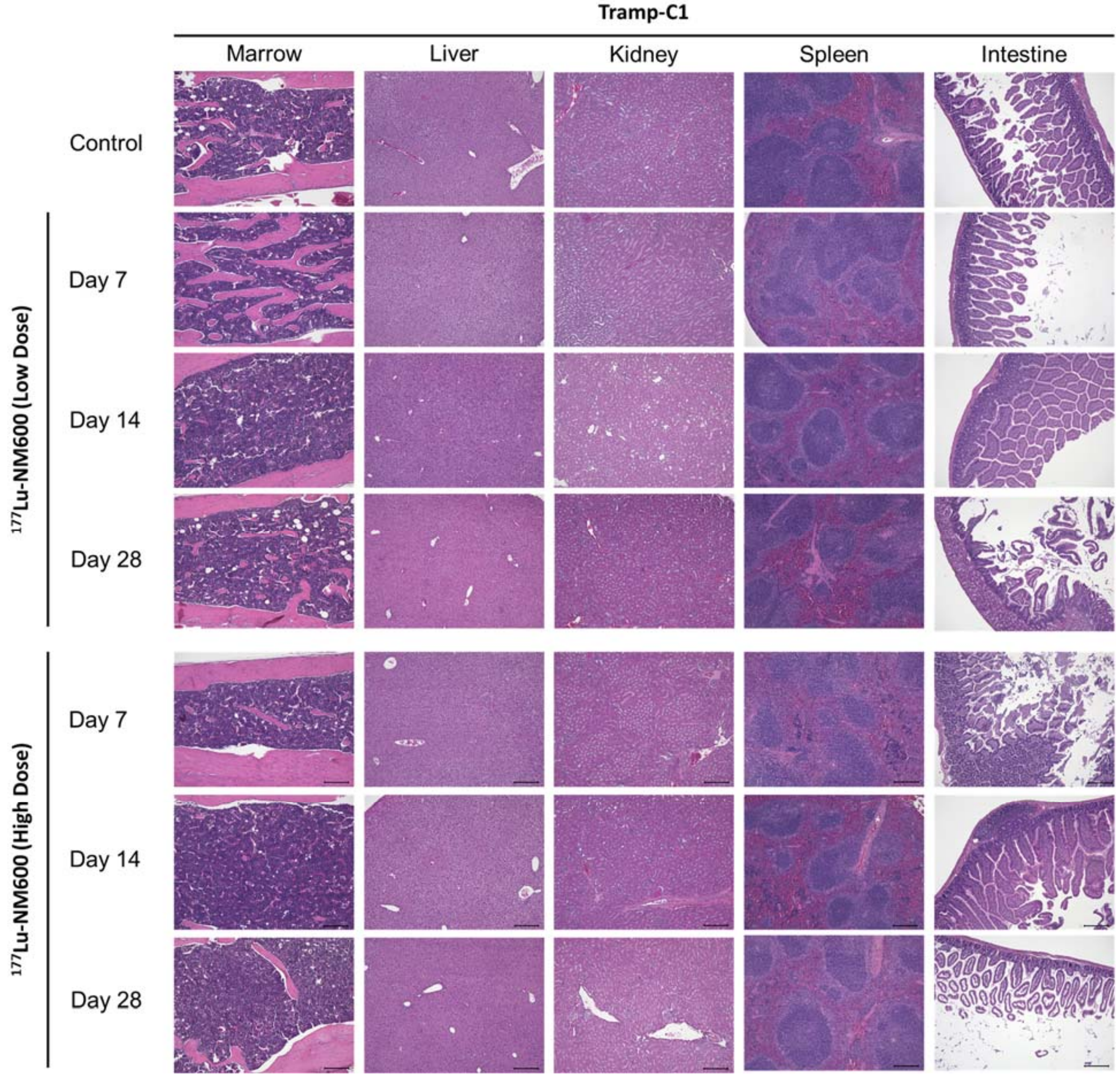


**Figure S7**. H&E staining of major organs collected 7, 14, or 28 days after administration of 5.55 MBq or 18.5 MBq of $^{177}$Lu-NM600 in Tramp-C1 tumor-bearing mice.

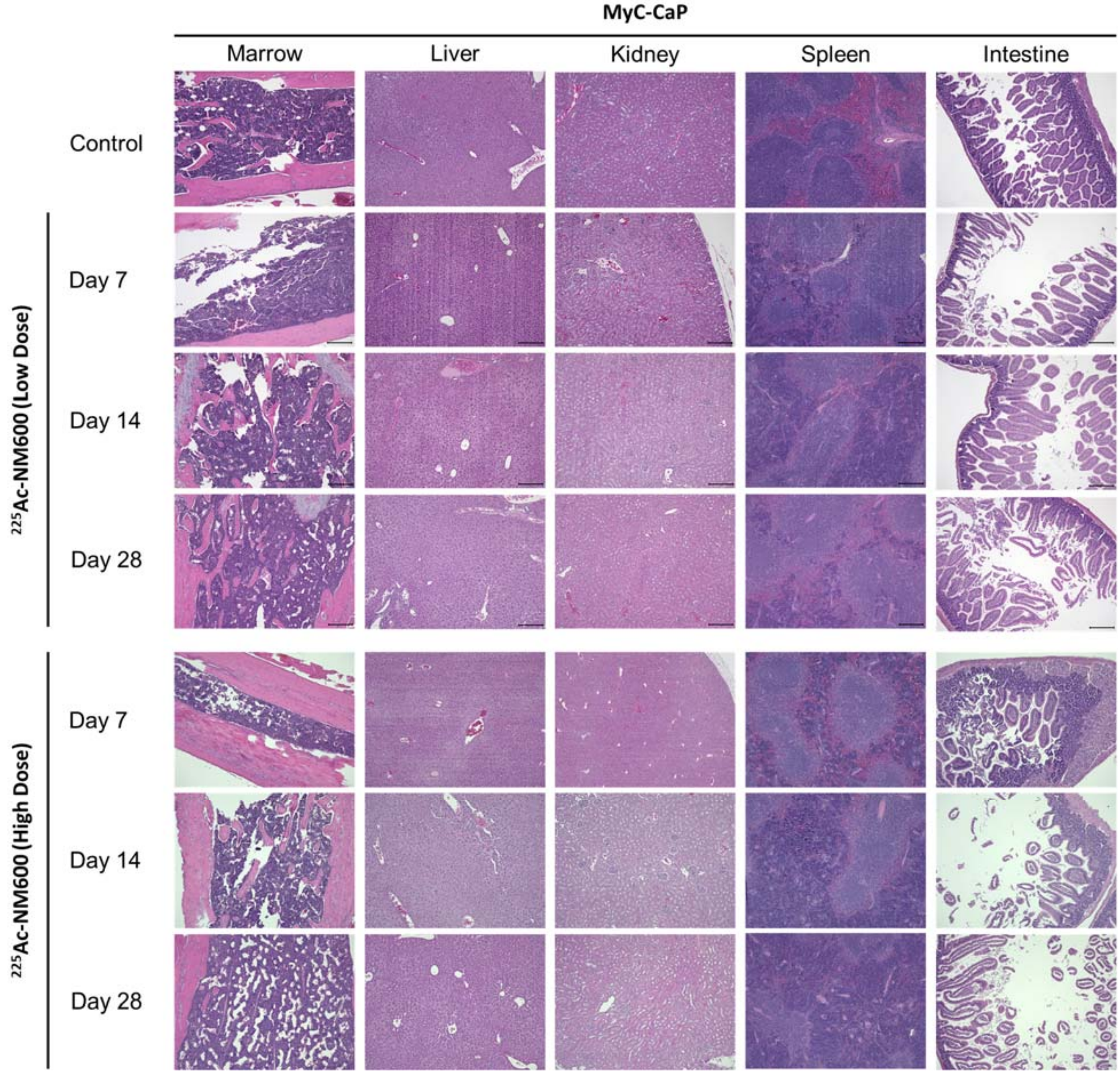


**Figure S8**. H&E staining of major organs collected 7, 14, or 28 days after administration of 7.4 kBq or 18.5 kBq of $^{225}$Ac-NM600 in Myc-CaP tumor-bearing mice.

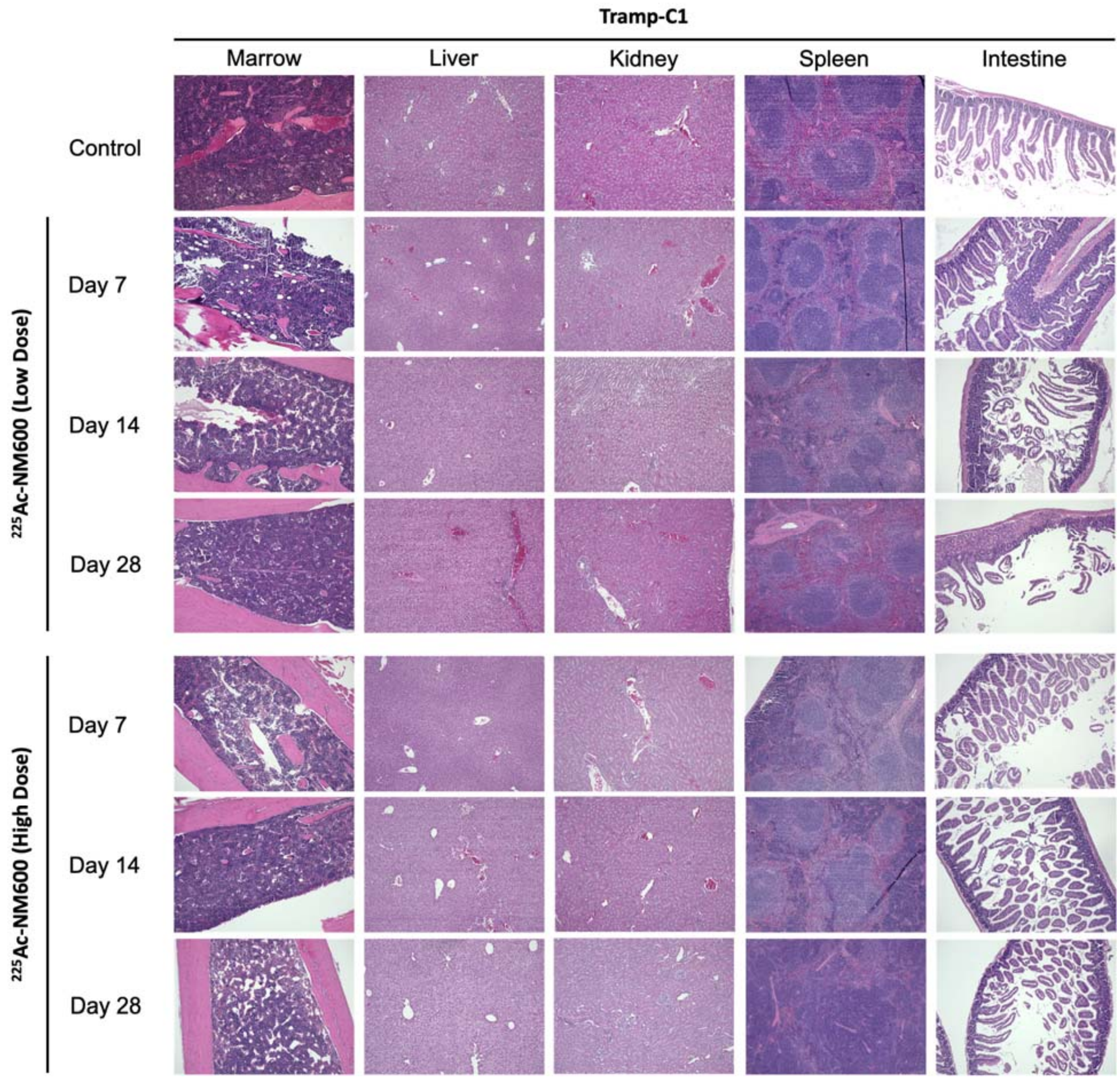


Figure S9. H&E staining of major organs collected 7, 14, or 28 days after administration of 7.4 kBq or 18.5 kBq of $^{225}$Ac-NM600 in Tramp-C1 tumor-bearing mice.

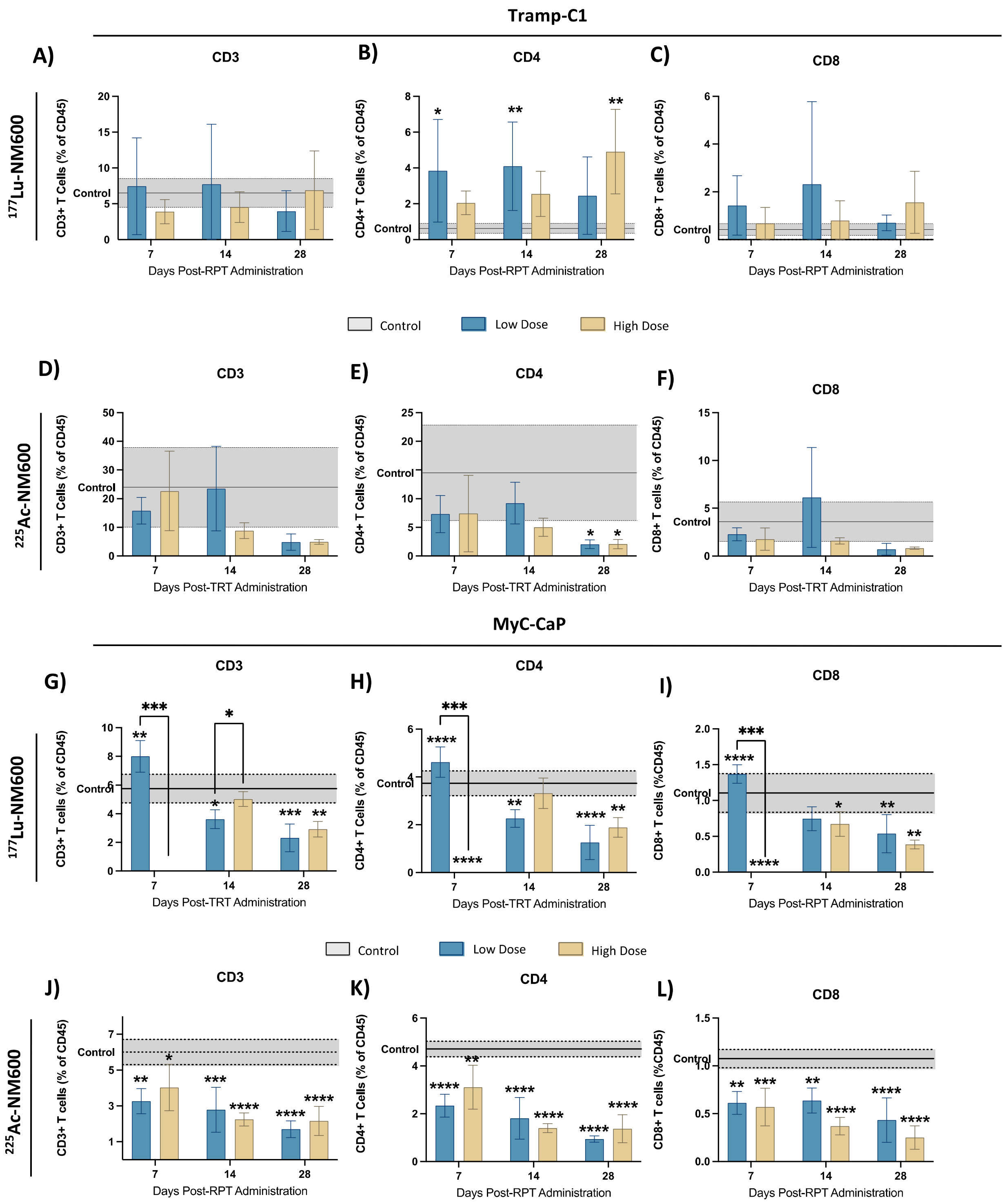


**Figure S10**. Flow Cytometry analysis of CD3, CD4, and CD8 markers in Tramp-C1 tumor-bearing mice that received A-C) $^{177}$Lu-NM600 or D-F) $^{225}$Ac-NM600 and in MyC-CaP tumor-bearing mice that received G-I) $^{177}$Lu-NM600 or J-L) $^{225}$Ac-NM600. Statistical analysis when compared to controls or otherwise noted. *$p<0.05$, ** $p<0.01$, *** $p<0.001$, ****$p<0.0001$

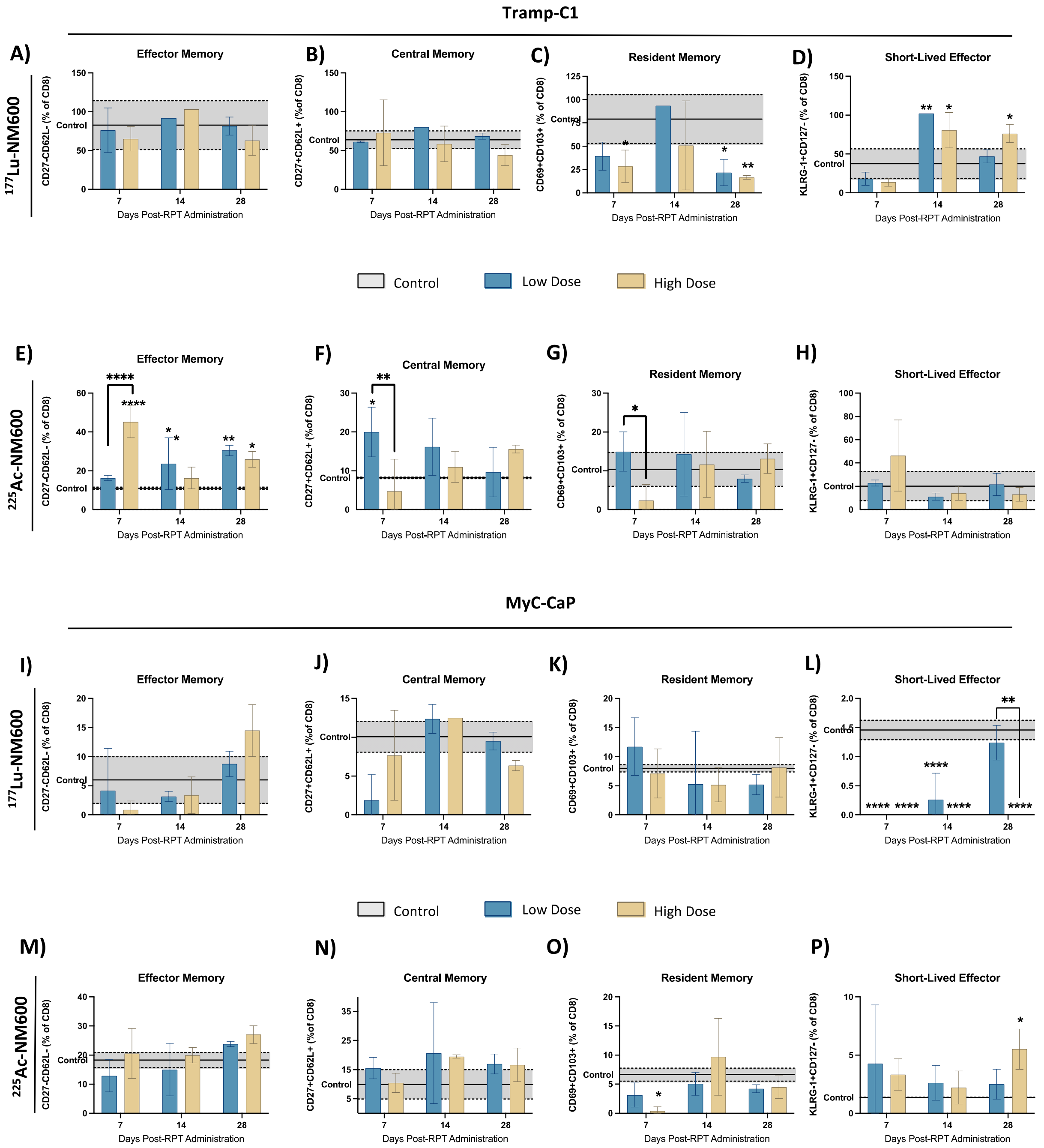


**Figure S11.** Expression levels of effector, central, resident, and short-lived effector memory markers in Tramp-C1 tumor-bearing mice that received A-D) $^{177}$Lu-NM600 or E-H) $^{225}$Ac-NM600 and in MyC-CaP tumor-bearing mice that received I-L) $^{177}$Lu-NM600 or M-P) $^{225}$Ac-NM600. Statistical analysis when compared to controls or otherwise noted. *p<0.05, ** p<0.01, *** p<0.001, ****p<0.0001.

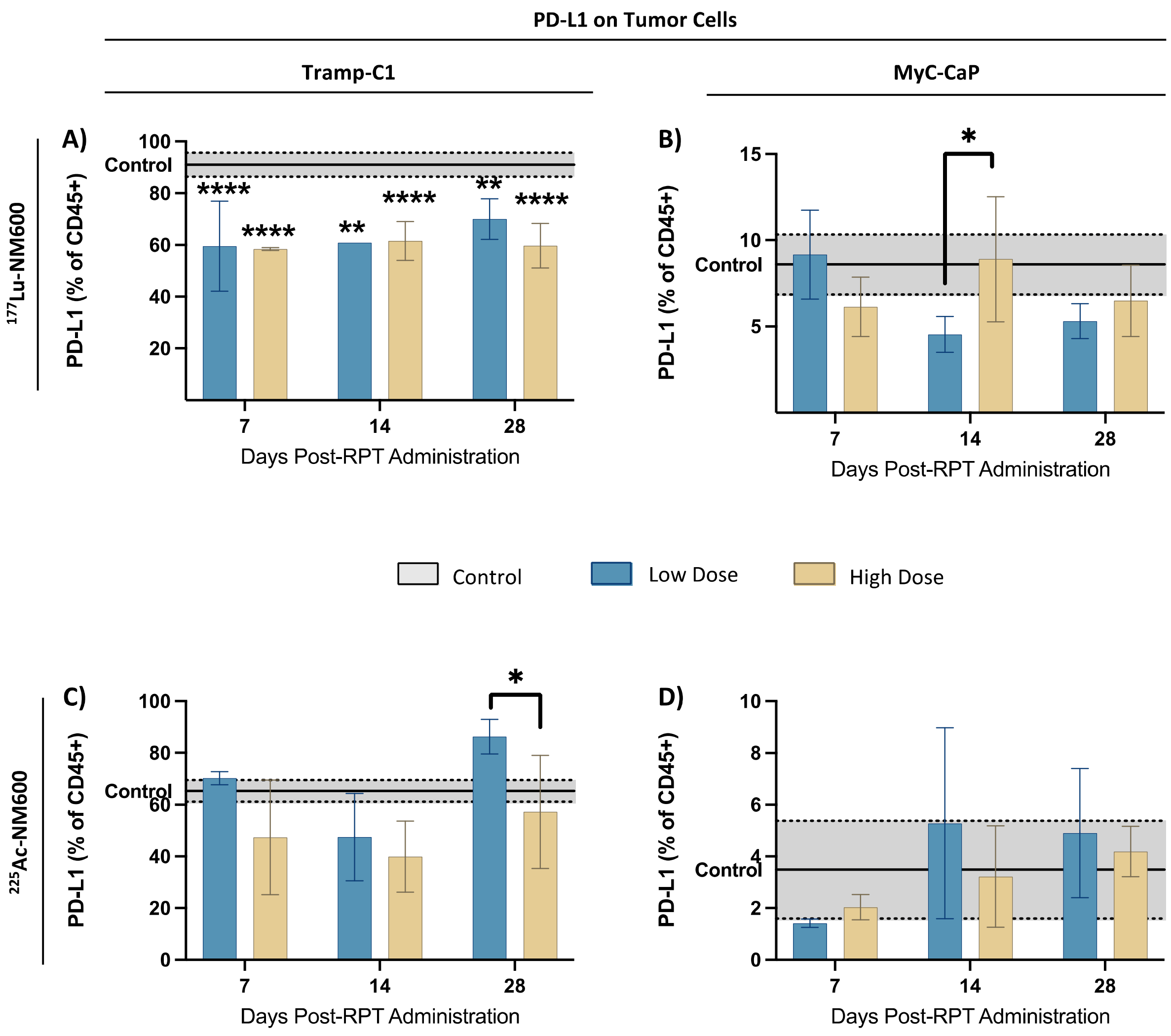


**Figure S12**. Flow Cytometry analysis of PD-L1 expression levels after administration of $^{177}$Lu-NM600 on A) Tramp-C1 and B) MyC-CaP tumors and after administration of $^{225}$Ac-NM600 on C) Tramp-C1 and D) MyC-CaP tumors (expressed as a percentage of CD45-). Statistical analysis when compared to controls or otherwise noted. *p<0.05, ** p<0.01, *** p<0.001, ****p<0.0001

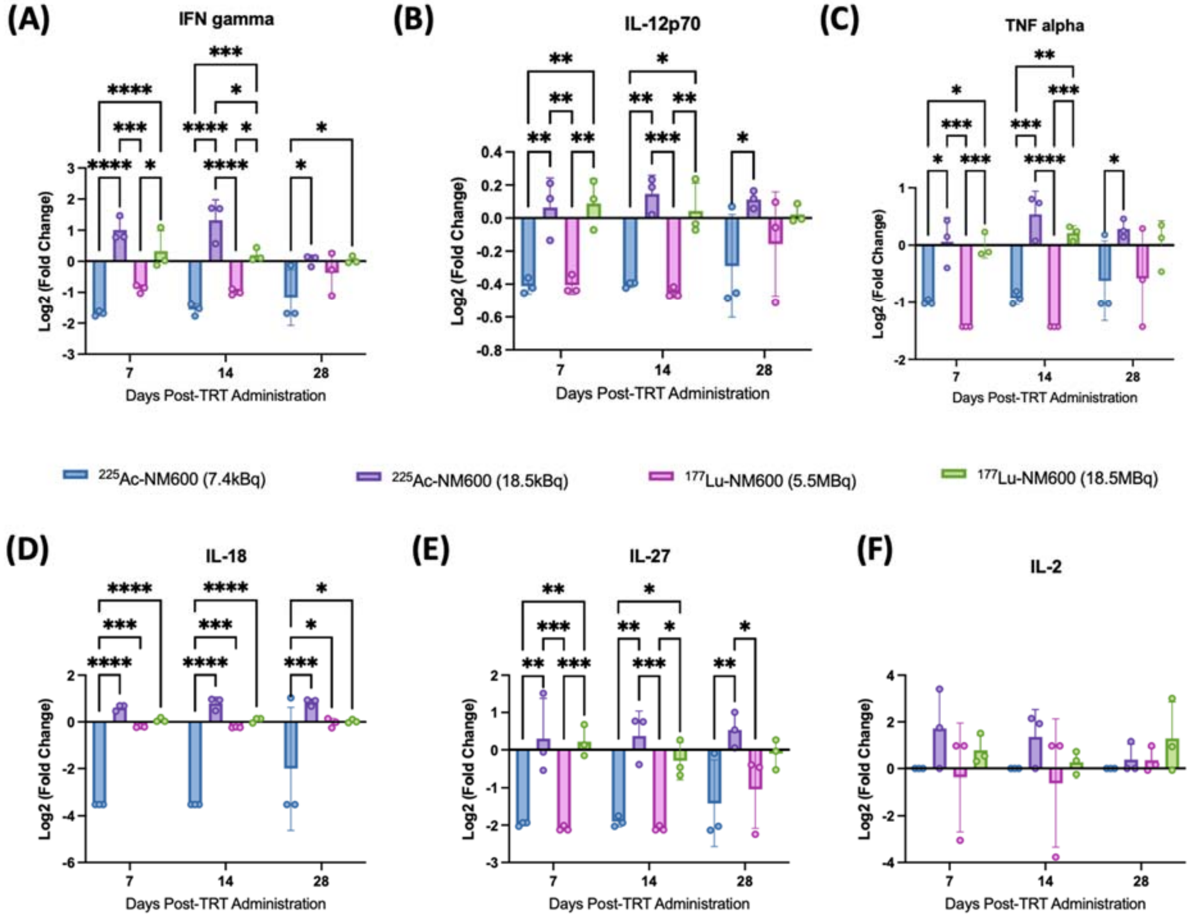


**Figure S13**. Analysis of the Th1 phenotype cytokines A) IFN gamma, B) IL-12p70, C) TNF alpha, D) IL-18, E) IL-27, F) IL-2 in TrampC-1 tumor-bearing mice after administration of .55 MBq or 18.5 MBq of $^{177}$Lu-NM600 or 7.4 kBq or 18.5 kBq of $^{225}$Ac-NM600. *p<0.05, ** p<0.01, *** p<0.001, ****p<0.0001.

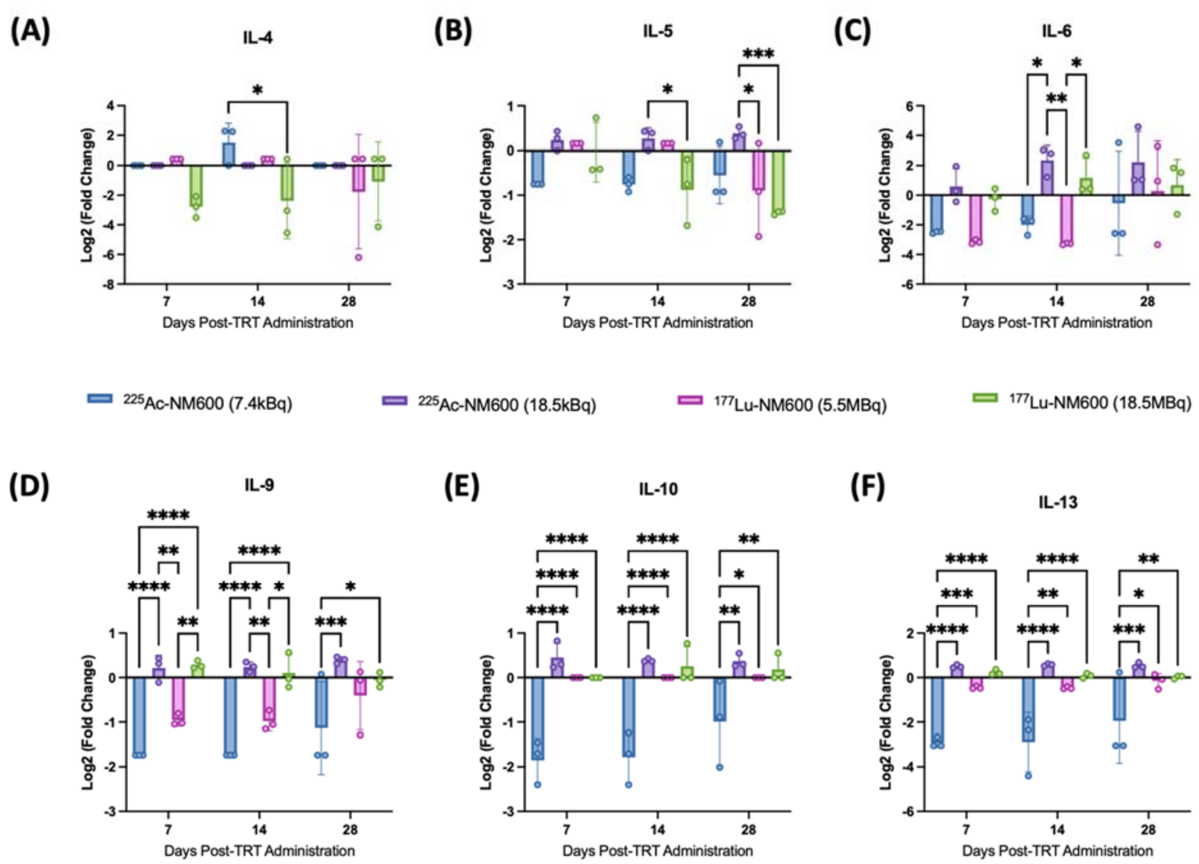


**Figure S14**. Analysis of the Th2 phenotype cytokines **A)** IL-4, **B)** IL-5, **C)** IL-6, **D)** IL-9, **E)** IL-10, and **F)** IL-13 in TrampC-1 tumor-bearing mice after administration of .55 MBq or 18.5 MBq of $^{177}$Lu-NM600 or 7.4 kBq or 18.5 kBq of $^{225}$Ac-NM600. *p<0.05, ** p<0.01, *** p<0.001, ****p<0.0001.

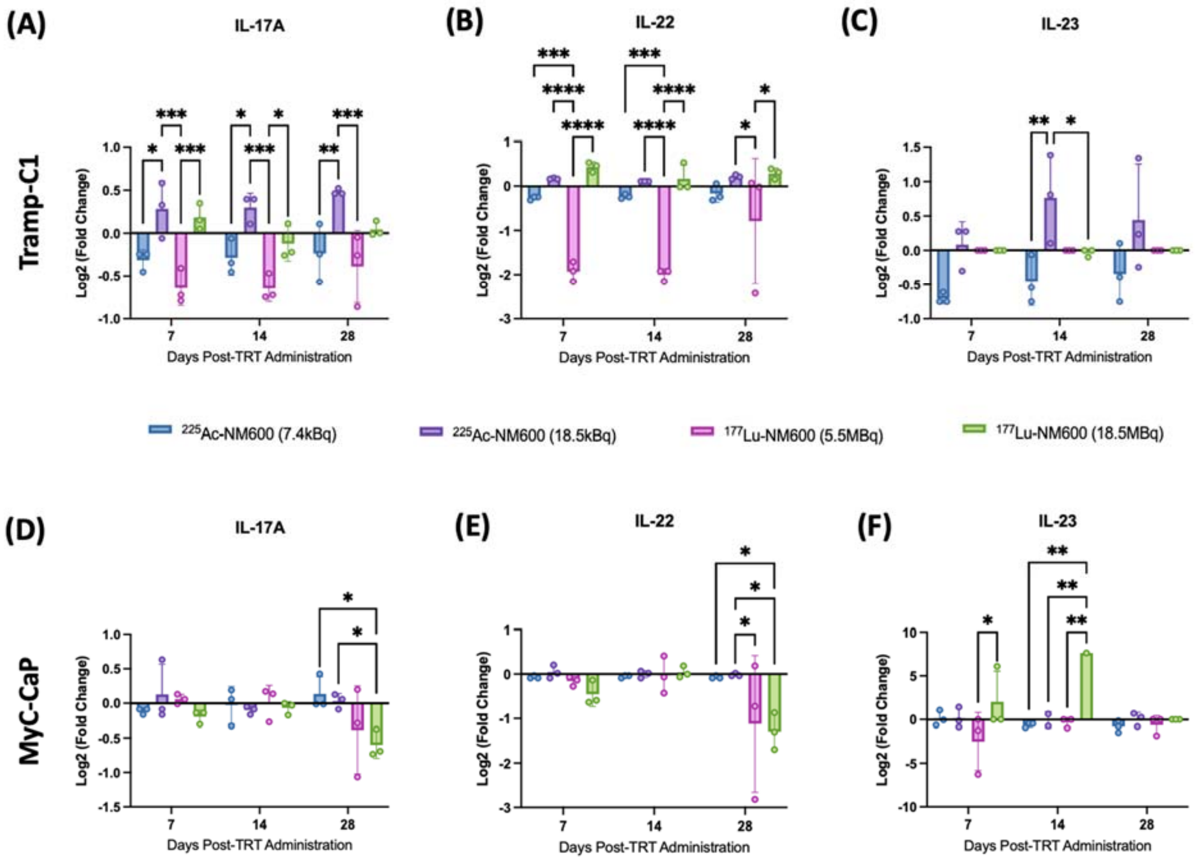


**Figure S15.** Analysis of the Th2 phenotype cytokines **A)** IL-17-A, **B)** IL-22 and **C)** IL-23 in TrampC-1 tumor-bearing mice and **D)** IL-17A, **E)** IL-22, **F)** IL-23 after administration of .55 MBq or 18.5 MBq of $^{177}$Lu-NM600 or 7.4 kBq or 18.5 kBq of $^{225}$Ac-NM600. *p<0.05, ** p<0.01, *** p<0.001, ****p<0.0001.

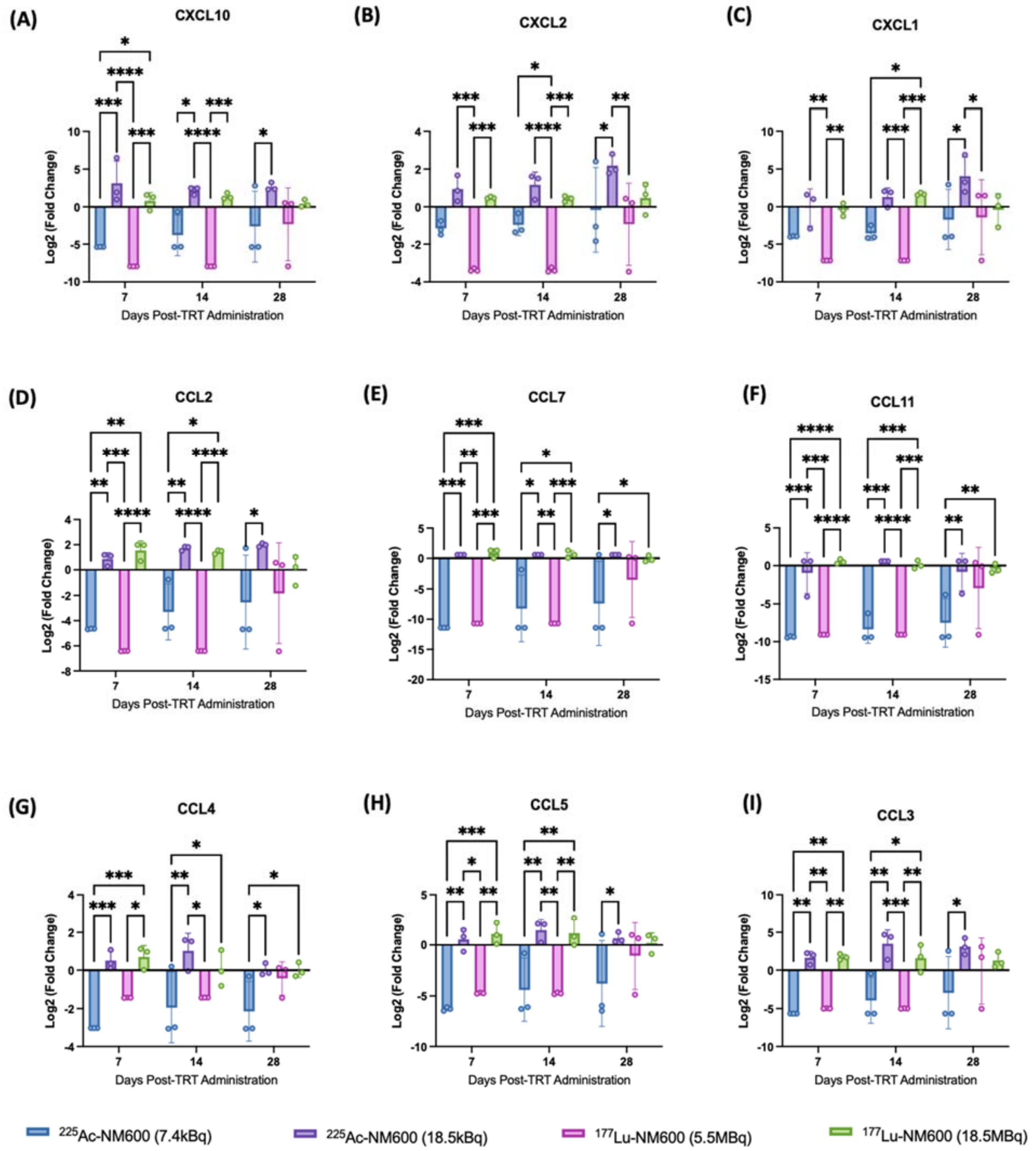


**Figure S16.** Analysis of **A)** CXCL10, **B)** CXCL2, **C)** CXCL1, **D)** CCL2, **E)** CCL7, **F)** CCL11, **G)** CCL4, **H)** CCL5, **I)** CCL3 chemokines in TrampC-1 tumor-bearing mice after administration of .55 MBq or 18.5 MBq of $^{177}$Lu-NM600 or 7.4 kBq or 18.5 kBq of $^{225}$Ac-NM600. *p<0.05, ** p<0.01, *** p<0.001, ****p<0.0001.

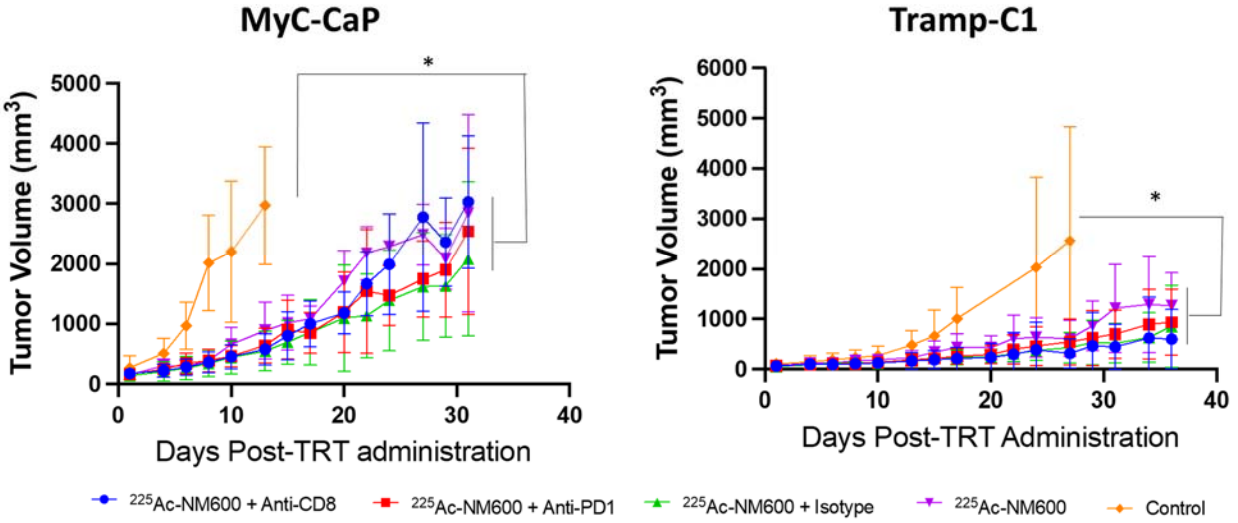


**Figure S17**. Tumor Growth Curves of MyC-CaP and Tramp-C1 tumor-bearing mice that received vehicle (control), 18.5 kBq of $^{225}$Ac-NM600, 18.5 kBq of $^{225}$Ac-NM600 + isotype, 18.5 kBq of $^{225}$Ac-NM600 + anti-PD1 (three doses on days 4, 7, and 10) or 18.5 kBq of $^{225}$Ac-NM600 + anti-CD8 (twice a week for 35 days). Control groups in both tumor models had significantly (*$p < 0.05$) higher tumor volumes when compared to any other group investigated.
